# Algorithmic Geometry Shapes the Biological Meaning of Thalamocortical Structural Covariance

**DOI:** 10.64898/2026.10.27.762356

**Authors:** Yuhan Liu, Valerie J. Sydnor, Alyssa Dai, Justine Ziolkowski, Shana Fu, Gabriel A. Devenyi, Theodore D. Satterthwaite, M. Mallar Chakravarty, Yohan Yee

**Affiliations:** Computational Brain Anatomy Laboratory, Cerebral Imaging Center, Douglas Mental Health University Institute, Montreal, QC, Canada; Experimental Imaging Center, Cumming School of Medicine, University of Calgary, Calgary, AB, Canada; Department of Psychiatry, University of Pittsburgh Medical Center, University of Pittsburgh, Pittsburgh, PA, USA; Center for the Neural Basis of Cognition, University of Pittsburgh and Carnegie Mellon University, Pittsburgh, PA, USA; McConnell Brain Imaging Centre, The Neuro (Montreal Neurological Institute–Hospital), Montreal, QC, Canada; B.Sc. Program in Neuroscience, Faculty of Science, University of Calgary, Calgary, AB, Canada; Department of Psychiatry, McGill University, Montreal, QC, Canada; Penn Lifespan Informatics and Neuroimaging Center, Perelman School of Medicine, University of Pennsylvania, Philadelphia, PA, USA; Department of Psychiatry, Perelman School of Medicine, University of Pennsylvania, Philadelphia, PA, USA; Center for Biomedical Image Computing and Analytics, University of Pennsylvania, Philadelphia, PA, USA; Department of Biological and Biomedical Engineering, McGill University, Montreal, QC, Canada; Department of Radiology, Cumming School of Medicine, University of Calgary, Calgary, AB, Canada

**Keywords:** structural covariance, thalamus, cortex, dimensionality reduction, non-negative matrix factorization, clustering, tractography, imaging transcriptomics

## Abstract

Neuroimaging studies often derive and discuss structural covariance (SC) as a representation of underlying brain circuitry that reflects coordinated maturation, anatomical connectivity, transcriptional similarity, and shared genetic influences. These associations motivate its use to study brain organization across development, aging, and disease. However, the algorithms used to derive SC networks impose constraints that may shape their biological interpretation. To understand the impact of algorithm choice, we used the thalamocortical system as an example and compared five common representations of voxelwise SC, namely, principal component analysis (PCA), independent component analysis (ICA), non-negative matrix factorization (NMF), orthogonal projective NMF (OPNMF), and *k*-means clustering, in 755 participants aged 8–22 years from the Philadelphia Neurodevelopmental Cohort (PNC). SC networks derived from deformation-based morphometry (DBM) were evaluated against three complementary biological benchmarks: age-related individual differences, fractional-anisotropy (FA)-based microstructural connectivity of regionally-specific thalamocortical connections, and Allen Human Brain Atlas gene expression. Each algorithm demonstrated underlying sensitivity to differing aspects of neurobiology: PCA and OPNMF showed the strongest associations between participant-level component weights and age. NMF, OPNMF, and *k*-means showed greater spatial overlap between positive cortical SC and tractography-derived projections. ICA and PCA showed stronger correspondence between thalamic and cortical gene-expression profiles across the transcriptome, whereas NMF, OPNMF, and *k*-means more often identified thalamic components with relatively high core-matrix gene expression. This study emphasizes that algorithmic constraints effectively act as biological priors and decomposition methods do not provide neutral compression of the data. We conclude that it is critical to consider whether the algorithmic objective matches the representational geometry of the neurobiology being studied. Overall, our findings suggest that SC studies should select and justify algorithms according to the biological question, explicitly considering whether their constraints favour distributed gradients, overlapping regional parts, or discrete clusters. Meanwhile, interpretations of SC findings could also be informed by the specific algorithmic constraints used to derive them.

## 1 Introduction

Structural covariance (SC; sometimes also referred to as anatomical covariance) refers to group-level statistical relationships between morphological measurements derived from magnetic resonance imaging (MRI) at different brain locations and estimated across participants (Mechelli et al., 2005; Carmon et al., 2020). Covariance is hypothesized to emerge from common developmental growth trajectories, shared genetic or environmental influences, activity-dependent plasticity, and direct or indirect anatomical coupling (Alexander-Bloch et al., 2013b; Schmitt et al., 2008; Bogado Lopes et al., 2023; Soriano-Mas et al., 2013; Carmon et al., 2020; Zhang et al., 2025; Kalantar-Hormozi et al., 2023). As one of the earliest studies using this technique, the Mapping Anatomical Correlations Across Cerebral Cortex (MACACC) study demonstrated that patterns of cross-sectional correlations in cortical thickness provided qualitative patterns that resembled tractography-derived connectivity maps (Lerch et al., 2006). In longitudinal developmental data, correlated rates of cortical thickness change have exhibited stronger maturational coupling within structurally and functionally interconnected cortical systems (Raznahan et al., 2011). Overall, studies have linked SC to coordinated maturation, anatomical connectivity, transcriptional similarity, and shared genetic influences (Mechelli et al., 2005; Alexander-Bloch et al., 2013b,a; Yee et al., 2018; Romero-Garcia et al., 2018). These findings point to SC being biologically informative but not mechanistically specific.

The biological interpretation of SC depends on how it is measured and represented. SC studies have used cortical thickness, surface area, volume, and tissue density (Mechelli et al., 2005; Carmon et al., 2020; Kalantar-Hormozi et al., 2023). Critically, as the structural features studied become more fine-grained, the enormous amount of pairwise relationships between each defined feature makes it difficult to characterize their overall covariance profile. The covariance relationships underlying these complex data can be uncovered by machine learning algorithms that perform dimensionality reduction, parts-based learning, and clustering. These approaches differ in how they weight the features in the data matrix and their specific constraints, e.g., whether they allow overlapping components or impose mutually exclusive assignments (Sotiras et al., 2015), directly shaping the resultant SC networks. Subsequently, SC networks can be represented as distributed gradients, spatially specific parts, or discrete groups of structures (Kharabian Masouleh et al., 2020; Sotiras et al., 2015; Chen et al., 2008). This suggest that different algorithms that use the same stuctural measures as input can yield representations that may emphasize different spatial relationships. Overall, this raises a crucial question that we will examine through the course of this manuscript; namely: Do SC networks identified using different algorithms provide varying, but meaningful, biological associations?

## 1.1 The thalamus and the cortex as a neuroanatomical system for assessing the sensitivity of different SC methods

Addressing this question comprehensively requires a systematic comparison between commonly employed algorithms. Also, they should be applied to a target brain system with well-defined anatomy and neurobiology which can be meaningfully represented by both continuous feature variation and discrete networks.

The thalamocortical system is ideal to test relationship between algorithmic choice and the relationship to the anatomy that it represents. Anatomically, classical thalamic nuclei have preferential and often reciprocal cortical projections, while molecular and connectivity properties also vary continuously across nuclear boundaries (Halassa and Sherman, 2019). Both focal sensorimotor and diffuse association-cortical connectivity patterns within the thalamus were found using tractography (Howell et al., 2024). Multimodal MRI findings identified thalamocortical structural connectivity gradients corresponding to intrathalamus microstructure, functional connectivity, and SC (John et al., 2025). A medial-to-lateral thalamic gene expression axis associated with anterior-to-posterior cortical connectivity was also observed using transcriptomics and diffusion tractography (Oldham and Ball, 2023). From the perspective of development, a recently published study used a thalamocortical tractography atlas to characterize a sensorimotor-association axis of maturation throughout childhood and adolescence (Sydnor et al., 2025). On the transcriptomic level, the relative expression of the calcium-binding proteins parvalbumin (*PVALB*) and calbindin (*CALB1* ) has been used to estimate core-rich and matrix-rich thalamus territories, which notably differ in cortical projection targets, with related molecular motifs recurring across multiple thalamic pathways (Phillips et al., 2019; Müller et al., 2020).

Given this organizational complexity across biological scales, a method that favours discrete territories may therefore capture one valid aspect of thalamic organization while obscuring another that is better expressed as a signed gradient, and vice versa. Leveraging the extensive resources on multiple aspects of thalamocortical relationships, our study compares the impact of algorithmic choices on SC networks on four fronts: their overlap with classical thalamic nuclei and cortical regions, how much they reflect age and sex, their relationship with tractography-derived connectivity, and their correspondence with transcriptomic information.

### 1.2 Deformation-based morphometry

For the present study, we sought a single morphological measure applicable to both the thalamus and cortex at voxelwise resolution. Deformation-based morphometry (DBM) quantifies local volumetric differences relative to a population template using Jacobian determinants derived from nonlinear image registration (Chung et al., 2001; Germann et al., 2025). Deriving these measurements from image deformation rather than tissue classification provides a tissue-agnostic approach for analysing both structures in a common volumetric space. Voxelwise sampling also allows examining fine-grained variation within the thalamus and cortex without averaging measurements within predefined nuclei or cortical parcels. This framework provides a common morphometric basis for comparing how different algorithms represent thalamocortical SC.

### 1.3 Decomposition algorithms

We examined independent component analysis (ICA), principal component analysis (PCA), non-negative matrix factorization (NMF), orthogonal projective NMF (OPNMF) and *k*-means clustering (*k*-means) as decomposition algorithm choices in deriving SC networks. All five approaches have been used to characterize structural covariance patterns across neurodevelopment (Sotiras et al., 2017; Sandini et al., 2018; Kalantar-Hormozi et al., 2023), healthy aging and cognition (Sotiras et al., 2015; Eckert et al., 2010; Wang et al., 2025), and neuropsychiatric and neurological disease (Xu et al., 2009; Abela et al., 2015; Sanfelici et al., 2022; Guo et al., 2023; Zheng et al., 2024; Robert et al., 2022; Patel et al., 2020). These algorithms impose markedly different representational properties (Table 1): PCA identifies orthogonal signed axes ordered by explained variance. ICA seeks statistically independent, usually non-Gaussian sources (Xu et al., 2009). NMF approximates a non-negative matrix with additive non-negative factors, often yielding localized, partly overlapping “parts” (Lee and Seung, 1999; Sotiras et al., 2015). OPNMF combines a projective reconstruction with non-negativity and orthonormality, increasing competition among components and promoting sparse, spatially segregated solutions (Yang and Oja, 2010; Wen et al., 2023). *K* -means replaces loading-based representation with hard assignment: each observation belongs to one nearest-centroid cluster. These objectives effectively determine which latent covariance patterns in the data are concentrated, separated, or suppressed, which may affect the specific biological information embedded in them.

**Table 1.** Mathematical and representational properties of the five methods.

| Method | Spatial weights / assignment | Typical map structure | Objective and constraints | Interpretation |
| --- | --- | --- | --- | --- |
| ICA | Signed continuous weights; voxels can contribute to more than one component. | Spatially independent component maps that may be focal or distributed and can overlap. | For whitened data $\mathbf{Z}$,<br>$\min_{\mathbf{R}} I(s_1, \dots, s_K)$<br>s.t. $\mathbf{S} = \mathbf{R}^\top \mathbf{Z}$, $\mathbf{R}^\top \mathbf{R} = \mathbf{I}_K$<br>(Hyvärinen and Oja, 2000) | Finds components that are as statistically independent as possible. Component order and the overall sign of each component are arbitrary (McKeown et al., 1998; Xu et al., 2009). |
| PCA | Signed continuous weights; voxels can contribute to more than one component. | Orthogonal component maps, often distributed across many voxels. | $\min_{\mathbf{U}} \ \mathbf{X} - \mathbf{U}\mathbf{U}^\top \mathbf{X}\ _F^2$ (Jolliffe and Cadima, 2016; Cunningham and Ghahramani, 2015)<br>s.t. $\mathbf{U}^\top \mathbf{U} = \mathbf{I}_K$ | Finds orthogonal directions that successively explain the greatest variance. Components are ordered by explained variance; the overall sign is arbitrary, while positive and negative weights indicate opposite directions along the same component (Friston et al., 1993). |
| NMF | Non-negative continuous weights in the factorized representation; features can contribute to more than one component. | Parts-based component maps that are often sparse or localized; overlap between components is allowed. | $\min_{\mathbf{W}, \mathbf{H}} \frac{1}{2} \ \mathbf{X} - \mathbf{W}\mathbf{H}\ _F^2$ (Lee and Seung, 2001; Lin, 2007)<br>s.t. $\mathbf{W} \geq 0$ , $\mathbf{H} \geq 0$ | Represents each observation as an additive combination of non-negative components. Because the factors are non-negative, components combine by addition rather than cancellation between positive and negative contributions (Lee and Seung, 1999; Sotiras et al., 2015). |
| OPNMF | Non-negative continuous weights in the factorized representation; exact non-negative orthogonality gives each input feature non-zero weight in at most one component. | Sparse, non-overlapping component maps with continuous variation within each component. | $\min_{\mathbf{W}} \ \mathbf{X} - \mathbf{W}\mathbf{W}^\top \mathbf{X}\ _F^2$ (Yang and Oja, 2010)<br>s.t. $\mathbf{W} \geq 0$ , $\mathbf{W}^\top \mathbf{W} = \mathbf{I}_K$ | Combines a projective non-negative representation with orthogonality, favoring sparse and complementary component maps with minimal overlap (Sotiras et al., 2015). |
| $k$-means Clustering | Hard one-of-$K$ assignment; each voxel belongs to exactly one cluster. | Discrete clusters; spatial contiguity is not imposed by standard $k$-means. | $\min_{\mathbf{G}, \mathbf{M}} \ \mathbf{X} - \mathbf{G}\mathbf{M}\ _F^2$<br>s.t. $\mathbf{G} \in \{0, 1\}^{V \times K}$ , $\mathbf{G}\mathbf{1}_K = \mathbf{1}_V$<br>(MacQueen, 1967; Lloyd, 1982) | Groups voxels with similar feature profiles and represents each cluster by its centroid. Unlike the factorization methods, $k$-means does not retain continuous voxel weights within a cluster (Thirion et al., 2014). |

*Note.* Matrices are written with voxels in rows, $\mathbf{X} \in \mathbb{R}^{V \times N}$ , and $K$ denotes the model order; transpose-equivalent forms apply under the opposite convention. $\mathbf{U}$ and $\mathbf{W}$ contain spatial component weights, $\mathbf{H}$ contains component coefficients, and $\mathbf{G}$ and $\mathbf{M}$ contain one-hot cluster assignments and centroid profiles, respectively. PCA assumes centered input. For ICA, $\mathbf{Z}$ denotes the whitened rank- $K$ representation, $\mathbf{R}$ the ICA rotation, and $\mathbf{S}$ the recovered spatial sources; the displayed expression is written in the transpose-equivalent spatial-ICA orientation.
For NMF and OPNMF, $\mathbf{X}$ in the displayed objectives denotes the non-negative representation supplied to the factorization. In this study, signed values were separated into positive and negative channels before factorization. Consequently, the exact non-overlap property of OPNMF applies to these factorization features (voxel-sign channels). After positive and negative channels are recombined into signed voxel maps, a physical voxel may contribute to more than one component through different sign channels. Descriptions of map structure summarize properties encouraged by each method’s mathematical constraints and should not be interpreted as guarantees of anatomical organization.

### 1.4 This study

Here, we compared PCA, ICA, NMF, OPNMF, and *k*-means in the same voxelwise thalamocortical DBM data. The sample comprised 755 participants aged 8–22 years from the Philadelphia Neurodevelopmental Cohort (PNC) after quality control and study-specific inclusion criteria (Satterthwaite et al., 2014). We evaluated component anatomy, age associations within sex, correspondence with FA-weighted projection maps from the PennLINC thalamocortical tractography atlas (Sydnor et al., 2025), and correspondence with gene expression from the Allen Human Brain Atlas (AHBA)(Hawrylycz et al., 2012). Connectivity was analysed separately for positive and negative covariance, and transcriptomics was examined both across the whole transcriptome and for *PVALB*/*CALB1*. Our main question is whether algorithmic mathematical constraints select different forms and scales of biological organization, and what that selectivity implies for the interpretation of SC networks.

## 2 Methods

### 2.1 Overview

We compared principal component analysis (PCA), independent component analysis (ICA), non-negative matrix factorization (NMF), orthogonal projective NMF (OPNMF), and *k*-means clustering to examine how algorithm choice shapes thalamocortical structural covariance (SC) representations and their biological associations. The study included 755 participants (female = 434) aged 8–22 years from the Philadelphia Neurodevelopmental Cohort (PNC), a community-based study of brain development in children, adolescents, and young adults (Satterthwaite et al., 2014). Deformation-based morphometry (DBM) of T1-weighted MRI provided a common voxelwise measure of local morphology in the thalamus and cortex. The comparison comprised four complementary analyses (Figure 1).

**Figure 1.**
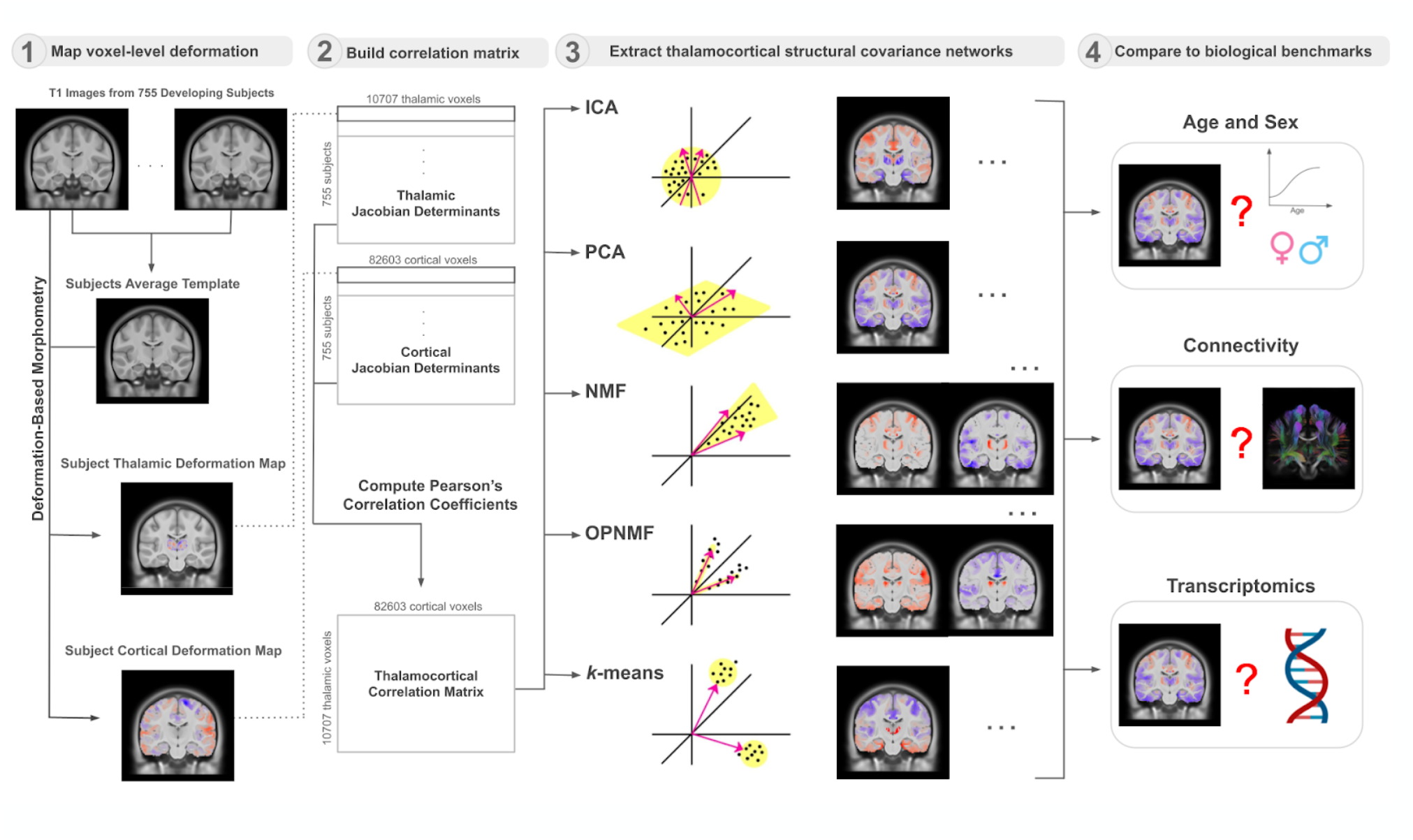
Overview of the analytical workflow used to derive voxelwise thalamocortical structural covariance networks and evaluate their correspondence with complementary developmental, anatomical, and transcriptomic measures.

(1) We characterized the spatial organization of group-level SC networks derived from each algorithm. Thalamocortical SC was estimated from correlations between thalamic and cortical DBM measurements across participants and SC networks were derived using each algorithm. We compared the resulting thalamic components and their associated cortical SC fields with atlas-defined anatomical regions to establish their localization and identify shared and method-dependent spatial patterns.

(2) In a separate decomposition of joint participant-level thalamic and cortical DBM measurements, we assessed each algorithm’s sensitivity to age-related individual differences. PCA, ICA, NMF, and OPNMF produced spatial components and weights describing their expression in individual participants. Associations between these weights and age were examined within each sex to compare the strength and direction of age associations across methods.

(3) We assessed the correspondence between group-level SC networks and anatomical connectivity. A reference thalamocortical tractography atlas was used to determine the strength of structural connectivity between the thalamus and individual cortical regions. For each thalamic component, a cortex-projected map of microstructural connectivity strength was associated to its cortical SC. We compared this projection map with the component’s associated cortical SC field to determine their spatial correspondence, evaluating positive and negative SC separately.

(4) We assessed the transcriptomic correspondence of group-level SC networks using the Allen Human Brain Atlas. We compared component-weighted thalamic and cortical gene-expression profiles to examine whether the paired SC patterns showed concordant or inverse molecular relationships. A complementary targeted analysis examined whether thalamic components emphasized territories with relatively high expression of *PVALB* or *CALB1*, distinguishing whole-transcriptome correspondence from localization of selected core-matrix-associated markers.

### 2.2 Dataset, image processing, and deformation-based morphometry

The PNC structural MRI release contained T1w images from 1,598 participants aged 8–22 years, acquired on a Siemens 3 T TIM Trio scanner using a 32-channel head coil and a magnetization-prepared rapid acquisition gradient-echo sequence (Satterthwaite et al., 2014). After image quality control using PyQC, conducted in accordance with previously described procedures (Bedford et al., 2020), participants were restricted to those with medical-history ratings of 0 or 1 and the required phenotypic data, yielding a final analytical sample of 755 participants. Acquisition parameters, exclusion counts, and quality-control procedures are provided in Supplementary Methods S1.

DBM quantifies local volumetric differences through the Jacobian determinants of image-registration transformations (Chung et al., 2001). We selected it to measure both anatomical structures within a common voxelwise framework. Images underwent visual motion quality control and intensity non-uniformity correction using iterative N4 processing (Tustison et al., 2010). Bias-corrected images were then processed with the population-template workflow described by Germann et al. (2025), using affine and nonlinear registration in Advanced Normalization Tools (ANTs) (Avants et al., 2008, 2011). Iterative template construction produced a study-specific anatomical reference from the preprocessed, registered participant images. Log-transformed relative Jacobian determinants indexed local morphological differences after excluding affine contributions to volume, reducing the influence of overall brain scaling.

Thalamic and cortical masks were defined using the Allen Human Reference Atlas–3D (AHRA) (Hawrylycz et al., 2012; Ding et al., 2016, 2020). We estimated a nonlinear transformation from the AHRA anatomical reference to the PNC population template and applied it to the atlas labels. Atlas alignment was visually inspected, and an additional registration-quality assessment excluded participant images with substantial misalignment within the thalamus or cortex. Details of template construction, atlas transformation, spatial sampling, and smoothing are provided in Supplementary Methods S1–S2.

Deformation fields from the final template-building iteration were used to generate relative Jacobian determinant maps. Relative Jacobian determinants exclude the volume contribution of the affine transformation and remove residual affine effects within the nonlinear deformation field (Germann et al., 2025). They were selected to emphasize regional volumetric variation after global affine normalization, reducing the contribution of overall brain scaling to estimates of thalamocortical structural covariance. The log-transformed relative Jacobians determinants were smoothed using Gaussian kernels with full widths at half maximum of 1, 2, and 4 mm. Positive and negative values indicate locally larger and smaller anatomy, respectively, relative to the population reference after affine adjustment. These maps provided spatially corresponding voxelwise indices of local volumetric variation across participants.

### 2.3 Preparation of structural covariance and participant-level inputs

At each voxel, log-transformed relative Jacobian determinants were residualized against the corresponding participant’s total thalamic or cortical volume. This structure-specific adjustment reduced the contribution of overall structure size to the local measurements.

For the spatial analyses, Pearson correlations were calculated across participants for every thalamic-cortical voxel pair. The resulting SC matrix, **R**, contained thalamic voxels in rows and cortical voxels in columns; each row described the cortical SC profile of one thalamic voxel. PCA, ICA, NMF, and OPNMF summarized variation among these profiles through graded voxelwise component weights, allowing continuous changes in component expression across the thalamus, albeit with different constraints on sign and spatial overlap. Such patterns could reflect gradient-like organization, although spatial smoothness was not explicitly enforced. In contrast, k-means assigned each voxel to a single cluster, discretizing variation among SC profiles. Each thalamic component or cluster was accompanied by a cortical coefficient field or mean SC profile, respectively.

For the age analyses, we retained the participant dimension by concatenating each participant’s thalamic measurements with spatially downsampled cortical measurements. The cortical voxels were downsampled to prevent cortical measurements from overpowering the number of thalamic measurements and dominating the age effects. This joint morphometric matrix, **Y**, was decomposed to obtain spatial patterns and their corresponding participant weights. These patterns were estimated separately from the components of **R**; the age analyses therefore assessed the algorithm families using a complementary input rather than testing age associations of the same group-level SC maps. *K* -means was excluded in this section because it consistently produces thalamus- and cortex-only clusters throughout different values of *k*.

PCA, ICA, and *k*-means accepted signed inputs. Because NMF and OPNMF only accept positive inputs, positive and sign-flipped negative entries of **R** were placed in separate cortical feature blocks and factorized together, preserving both correlation directions in a non-negative representation. In the positive block, negative entries were replaced by zero values, and vice versa. For the participant-level analyses, positive and sign-flipped negative entries of **Y** were factorized separately, with the same zero-out scheme. In this second pathway, the two solutions refer to the sign of the volume-adjusted morphometric measurements. Matrix definitions and sign handling are detailed in Supplementary Methods S3.

### 2.4 Algorithm objectives, implementation, and model order

Each method summarized the input as spatial components or clusters together with their associated feature coefficients. Following Table 1, **X** ∈ ℝ^*V* ×*N*^ denotes the input with *V* voxels in rows and *N* features in columns. Columns represented cortical features for the group-level SC analyses and participants for the age-sensitivity analysis. Here, *K* denotes the number of components or clusters, **I**_*K*_ is the *K* × *K* identity matrix, and 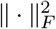 denotes the sum of squared matrix entries. For PCA, **X** denotes centred input; for NMF and OPNMF, it denotes the non-negative representation constructed using the sign-handling procedures described above. The objectives and constraints below correspond to Table 1; implementation settings are reported in Supplementary Methods S4.

- **ICA: statistically independent spatial sources**. ICA rotates the whitened data to minimize statistical dependence among the recovered sources (Hyvärinen and Oja, 2000):

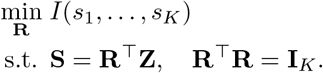

Here, *I*(*s*_1_, …, *s*_*K*_) denotes mutual information among the sources. This expression uses the spatial-ICA orientation: **Z** contains the centred, whitened rank-*K* representation with voxels in columns, and the rows of **S** contain the spatial sources. The symbol **R** denotes the ICA rotation in this objective, rather than the SC matrix defined earlier. FastICA pursued the independence criterion using a non-Gaussianity contrast (Hyvärinen and Oja, 1997, 2000). For the group-level SC analysis, the sources provided signed thalamic maps, and their cortical coefficients were projected back from the whitened space. Component order and overall sign were arbitrary (McKeown et al., 1998; Xu et al., 2009).
- **PCA: orthogonal axes ordered by explained variance**. PCA estimates an orthonormal spatial basis that minimized reconstruction error for the centred input (Jolliffe and Cadima, 2016; Cunningham and Ghahramani, 2015):

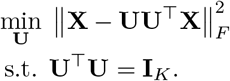

The columns of **U** define the spatial component directions, with corresponding feature coefficients given by **U**^⊤^**X**. For group-level SC, the representation paired thalamic spatial maps with cortical coefficient profiles. Principal components are ordered by explained variance and could overlap spatially. Positive and negative weights described opposing contributions within a component, while its overall sign was arbitrary (Friston et al., 1993; Jolliffe and Cadima, 2016).
- **NMF: additive components with overlapping membership**. NMF approximated the non-negative input using spatial weights **W** and feature coefficients **H** (Lee and Seung, 2001; Lin, 2007):

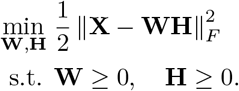

The columns of **W** contained spatial component weights, and the rows of **H** contained their associated feature coefficients. Non-negativity meant that components combined additively, while allowing each voxel to contribute to multiple components (Lee and Seung, 1999; Sotiras et al., 2015). For group-level SC, the cortical coefficients are separated after fitting into the positive- and negative-covariance networks associated with each thalamic component.
- **OPNMF: non-negative components with restricted overlap**. OPNMF combined a projective reconstruction with non-negativity and orthogonality (Yang and Oja, 2010):

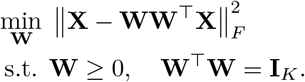

The columns of **W** contained spatial component weights, and the associated feature coefficients were given by **W**^⊤^**X**. Under exact non-negative orthogonality, each row of **W** can have non-zero weight in at most one component, restricting spatial overlap while retaining continuous weights within each component (Sotiras et al., 2015). The numerical implementation approximated the orthogonality constraint. Continuous weights were retained for quantitative analyses. As with NMF, the cortical coefficients from the group-level SC analysis retained separate positive- and negative-covariance networks.
- ***K* -means: mutually exclusive groups of similar profiles**. *K* -means grouped voxels by minimizing the reconstruction error between their feature profiles and their assigned cluster centroids (MacQueen, 1967; Lloyd, 1982):

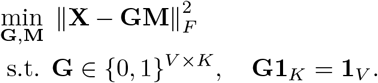

Here, **G** contains binary voxel-to-cluster assignments, **M** contains the cluster centroid profiles, and **1**_*K*_ and **1**_*V*_ are vectors of ones. The assignment constraint places each voxel in exactly one cluster. For group-level SC, the outputs were binary thalamic memberships and the mean signed cortical SC profile of each cluster. Cluster labels were arbitrary, and spatial contiguity was not imposed (Thirion et al., 2014).

PCA, ICA, NMF, and *k*-means were implemented using scikit-learn version 1.7.1 (Pedregosa et al., 2011), with custom whitening for ICA; OPNMF used the `juaml/opnmf` implementation. Candidate model orders ranged from *K* = 2 to 25. Assessment of component numbers used repeated hold-out analyses for the factorization methods and clustering diagnostics for *k*-means. For the purpose of comparison, we did not seek to establish individually optimal model order for every algorithm. The reported comparisons used a common *K* = 9 due to its balanced performance throughout algorithms. Implementation settings and model-order assessment are described in Supplementary Methods S4.

### 2.5 Anatomical characterization

We first examined where each algorithm placed its thalamic and cortical components and how these distributions relate to recognizable anatomical territories. Component maps were compared with individual AHRA thalamic and cortical labels.

Signed maps were separated into positive and negative-magnitude fields. Each nonempty field was divided by its maximum value, yielding weights between zero and one; non-negative NMF and OPNMF fields were normalized in the same way. Thalamic *k*-means memberships remained discrete. Anatomical correspondence was measured with soft Dice overlap, which compares continuous component weights with a binary atlas mask without first converting the component to a hard boundary. Ordinary Dice overlap was used for binary thalamic clusters from *k*-means. Larger scores indicated greater correspondence between the normalized component and the atlas territory.

For the component-by-region heatmaps, the larger of the positive- and negative-side overlap scores was retained separately for each component-region pair, with ties assigned to the positive side. Label selection, normalization, overlap definitions, and display rules are described in Supplementary Methods S5.

### 2.6 Age associations within sex

The developmental analysis asked whether the methods concentrated age-related individual differences into their participant-level components. This provides a complementary assessment of representations derived from a cohort spanning childhood, adolescence, and young adulthood, during which cortical morphology and thalamocortical connectivity exhibit notable age differences (Sotiras et al., 2017; Sydnor et al., 2025). We estimated age associations separately within females and males to examine their direction and consistency.

PCA, ICA, NMF, and OPNMF were fitted to the joint participant-level morphometric input described above. The resulting participant weights indexed the expression of each spatial pattern in individual participants. PCA and ICA weights were signed; NMF and OPNMF weights were non-negative and retained separately for the positive- and negative-value solutions. Under the joint-morphometry implementation used here, *k*-means consistently produced several clusters restricted to either the thalamus or cortex and was excluded from the primary comparison of paired thalamocortical participant weights.

For each component, participant weights were regressed on age within each sex. The signed *t* statistic for the age coefficient summarized the direction and statistical strength of the association; for PCA and ICA, its sign was interpreted relative to the retained spatial orientation. Pairwise correlations between participant-weight vectors within and across methods assessed whether different components represented similar individual differences. The regression specification and multiplicity correction are given in Supplementary Methods S6.

### 2.7 Correspondence with tractography-derived thalamocortical structural connectivity

The connectivity analysis asked whether cortical regions that exhibited high structural covariance with particular areas of the thalamus were also strongly structurally connected to those thalamic locations. To assess this, we used a validated thalamocortical tractography atlas comprising streamlines connecting the thalamus and cortex, together with microstructure-derived information about the connecting pathways (Sydnor et al., 2025). For each thalamic component, we identified its cortical projection territory using the atlas streamlines and compared the corresponding cortical SC field with the tractography-derived projection map, weighted by mean fractional anisotropy (FA) as an index of pathway microstructure. The reference was the PennLINC thalamocortical tractography atlas in HCP1065 1.25-mm space, a curated structural connectivity atlas comprising streamlines connecting thalamus and cortex (Sydnor et al., 2025). Thalamic and cortical SC component maps were transformed into this atlas space before comparison.

Continuous thalamic maps generated by each SC algorithm were converted to seed masks by retaining the highest-weight 15% of thalamic voxels after comparing thresholds of 10%, 15% and 20%; This selection identified the thalamic locations where each SC component was expressed most strongly. For PCA and ICA, upper- and lower-tail masks represented the positive and negative component poles. This threshold provided a common selection rule for continuous maps while limiting fragmentation; binary *k*-means clusters were used at their native sizes. Each thalamic mask was used to filter thalamocortical tractography atlas streamlines either by intersection with the mask or by an endpoint lying within it, producing component-specific cortical projection fields. Transformation and filtering details are provided in Supplementary Methods S7.

For the FA-weighted comparison, cortical projection masks were weighted using atlas-derived mean fractional anisotropy (FA) for the corresponding thalamocortical connections. These weights were assigned at the cortical-parcel level rather than estimated separately within each component’s filtered streamlines. The result was a cortical map where the values represent mean FA in the structural pathway connecting that cortical region to the thalamus. The reference therefore combined projection location with connection-level microstructural weighting, rather than measuring component-specific connection strength.

Cortical SC maps were separated by sign and normalized to [0, 1] within a common cortical mask. For PCA and ICA, same-sign thalamic and cortical poles defined positive component contributions, whereas opposite-sign poles defined negative contributions. For NMF, OPNMF, and *k*-means, the two directions were the positive and negative cortical fields associated with each thalamic component.

We evaluated three complementary outcomes: soft Dice overlap summarized overall spatial correspondence while retaining graded map weights; sensitivity summarized coverage of the reference projection field; and specificity summarized agreement outside that field. Component-level scores and their distributions across algorithms were examined separately for positive and negative SC. The exact map definitions, metric formulas, and aggregation across thalamic poles or filtering conditions are specified in Supplementary Methods S7.

### 2.8 Whole-transcriptome and thalamic cell type marker expression correspondence

The transcriptomic analyses addressed two questions: whether each paired thalamic and cortical SC pattern showed a relationship across many genes, and whether its thalamic component emphasized territories with relatively high expression of selected molecular markers. Gene-expression data were obtained from the Allen Human Brain Atlas (AHBA) (Hawrylycz et al., 2012). Thalamic and cortical samples were processed and normalized separately, with 17,205 genes retained after probe filtering and annotation. Refined sample localizations (Yee et al., 2025) were related to the PNC template to align expression measurements with the component maps. Processing, sample assignment, and weighting details are provided in Supplementary Methods S8.

For each component and analysed covariance direction, component-weighted mean expression was calculated separately in the thalamus and the corresponding cortical field. Continuous spatial weights were used for the factorization methods, and binary memberships defined the thalamic contribution for *k*-means. Whole-transcriptome correspondence was summarized by regressing cortical mean expression on thalamic mean expression across the retained genes and extracting the slope *t* statistic. Positive statistics indicated concordant profiles and negative statistics indicated inverse profiles. Where one value was displayed per component, the covariance direction with the larger absolute statistic was retained with its sign preserved.

The targeted analysis examined *PVALB* and *CALB1*, which encode parvalbumin and calbindin genes and have been used to characterize core-matrix-associated thalamic organization (Jones, 1998; Müller et al., 2020). This analysis tested marker localization within the thalamus rather than thalamus-cortex correspondence across the full transcriptome. For the dominant covariance direction of each component, thalamic component-weighted expression was standardized separately for each marker across all algorithm-component combinations. Positive *z* scores therefore identified relatively marker-rich components within the comparison set; they did not indicate enrichment against a spatial null or establish a distinct cellular population (Phillips et al., 2019).

## 3 Results

### 3.1 Algorithms recover spatially distinct thalamocortical SC networks with shared structures

To get an overview of the localization of the components, we compared the thalamic and cortical SC components from each algorithm to atlas-defined thalamic nuclei and cortical regions, as shown in the component-by-region heatmaps (Figure 2).

**Figure 2.**
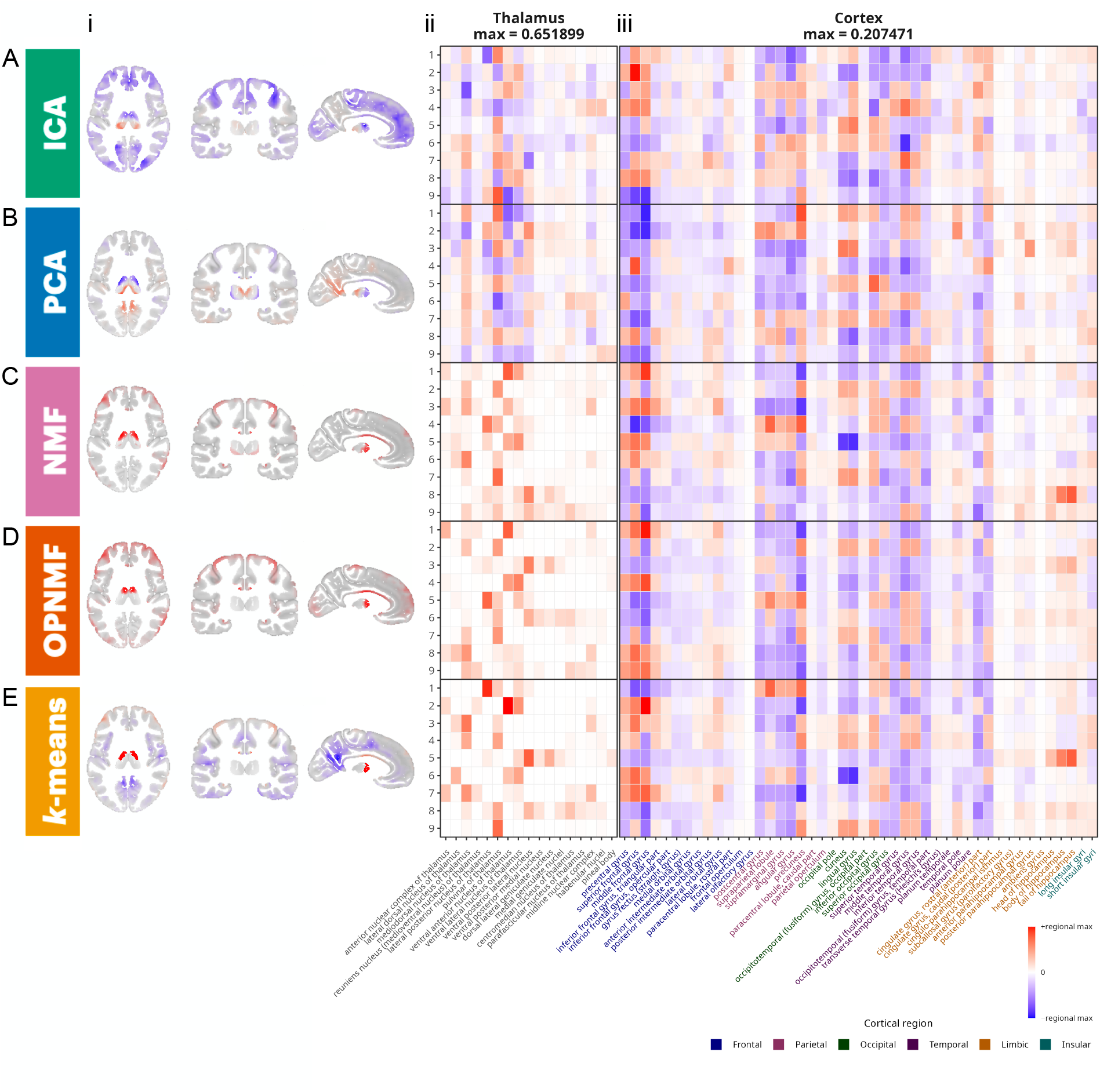
Spatial organization of thalamocortical structural covariance components across algorithms. **A–E**, results for ICA, PCA, NMF, OPNMF, and *k*-means, respectively. Within each algorithm, **(i)** sagittal, coronal, and axial views show the component with the closest spatial correspondence to PCA component 1; **(ii)** heat maps summarize overlap between thalamic component loadings and atlas-defined thalamic nuclei; and **(iii)** heat maps summarize overlap between cortical component loadings and cortical regions grouped by anatomical system or lobe. Signed values in PCA and ICA indicate opposing poles of the same component; their global sign is arbitrary. NMF and OPNMF maps are non-negative within each sign-separated representation, whereas *k*-means memberships are binary and mutually exclusive. Regional abbreviations should be defined in the final figure key.

All five methods recovered components with shared, recognizable thalamic and cortical anatomy, but they differed sharply in spatial representations (Figure 2). PCA and ICA produced broad signed components extending across several thalamic nuclei and distributed cortical territories. By contrast, NMF and OPNMF concentrated loadings into sparser thalamic and cortical fields, and *k*-means produced spatially exclusive clusters for the thalamus, each paired with a distributed cortical covariance profile. Despite these differences, a recurring anatomical pattern distinguished posterior-association covariance involving the pulvinar and mediodorsal thalamus with parietal, temporal, and occipital cortex, from anterior-frontal/sensorimotor covariance involving ventral anterior thalamus with frontal and precentral regions. Overall, these methods converged on elements of a common thalamocortical covariance architecture, while representing them differently as distributed graded components, localized and potentially overlapping parts, or discrete thalamic clusters. The first principal component (PC1) in PCA explains the largest percentage of variance and is often expressed in literature as the major gradient. Here, PC1 (42.23% variance explained) captures a posterior-association versus anterior-frontal/sensorimotor covariance: one pole loaded most strongly on the pulvinar and, to a lesser extent, the mediodorsal thalamus, together with the precuneus, middle and inferior temporal cortex, cuneus, and lingual gyrus. The opposing pole emphasized ventral anterior thalamus and the middle, precentral, and superior frontal gyri, with more modest involvement of the supramarginal gyrus (Figure 2B). Analogous components were found in other algorithms by pairing through Pearson’s correlations and shown as demonstrations (Figure 2i). Notably, the same axis was preserved but with minor variations in degrees of overlap with anatomical features. Even in these highly correlated components, there are sharp differences in the representations of these components and, to a minor extent, different magnitudes of overlap with anatomically defined structures.

These differences were quantified in detail through the heatmap of regional overlap (Figure 2). Some localized components were dominated by a single thalamic territory. For example, a *k*-means cluster centred on the lateral posterior nucleus; but most components crossed two or three nuclei, and several remained distributed across a larger portion of the thalamus. Thus, even the hardest partition did not simply reproduce a classical nuclear atlas. Across the cortex, the largest absolute overlap occurred repeatedly in precentral, superior frontal, and middle frontal gyri; broad parietal cortex; the cuneus and lingual gyrus; and, more moderately, temporal cortex. Middle temporal overlap was especially prominent in several ICA components. Cingulate regions showed moderate involvement, whereas olfactory territories were consistently near zero.

NMF (component 8) and *k*-means (cluster 5) each contained a pair of similar components with positive covariance in hippocampal body and tail, mild negative covariance in frontal and parietal cortex, and a thalamic field centered on the ventral posterolateral territory with weaker lateral dorsal and dorsolateral posterior involvement. These components show that localized methods did not merely isolate canonical thalamocortical sensory sectors; they could also identify compact limbic-associated covariance motifs.

### 3.2 PCA and OPNMF show the strongest associations with age

Distinctly, age effects differed more by algorithm than by sex, and were statistically largest in PCA and OPNMF (Figure 3A). PCA yielded seven significant age-associated components with absolute *t* values of approximately 3 to 6: four were detected in both sexes and three in females only. OPNMF produced the broadest age-related profile. Eight positive components were significant, with magnitudes similar to PCA: four were present in both sexes, two in females only, and two in males only. Seven negative OPNMF components also met the threshold: four in both sexes, two in males only, and one in females only. Significant components detected in both sexes generally showed concordant directions and among the largest effect magnitudes.

**Figure 3.**
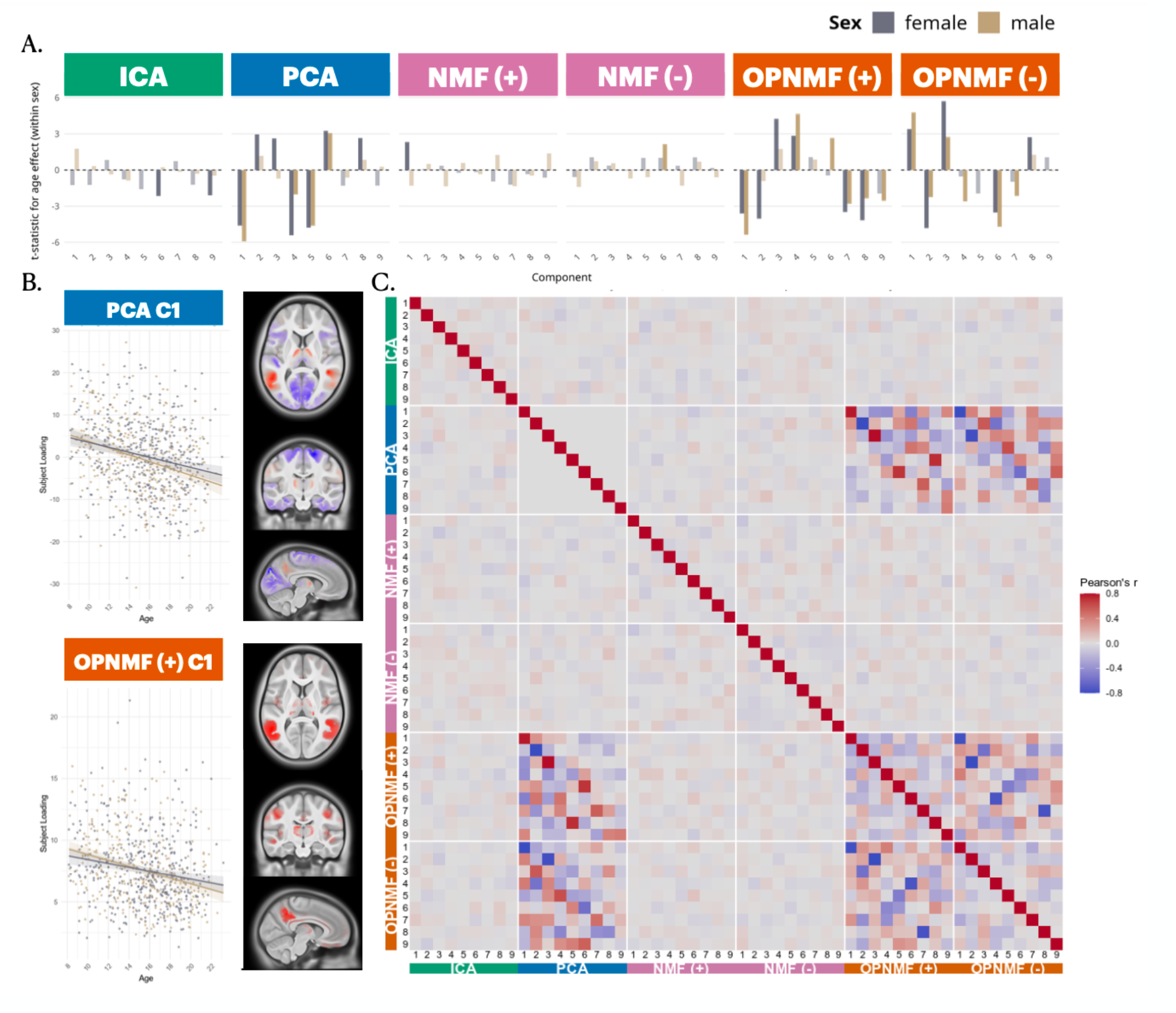
Age-related expression of thalamocortical structural covariance components. **A**, sex-stratified *t* statistics for the age term in component-loading models, shown separately for ICA, PCA, positive and negative NMF, and positive and negative OPNMF. Components that did not reach statistical significance are grayed. *k*-means is omitted because several clusters were confined to one anatomical structure and did not yield comparable paired thalamocortical subject loadings. **B**, representative age effects and spatial maps for **(i)** PCA component 1 and **(ii)** positive OPNMF component 1. Scatterplots show subject loadings against age with sex-specific linear fits; maps show the corresponding thalamic and cortical component fields. **C**, pairwise correlations among subject loadings for all components and algorithms. PCA within-method correlations are near zero by construction.

ICA yielded only two components that met the multiple-comparison threshold; both effects were observed in females and were modest in magnitude (absolute *t* values near 2). No positive NMF component met the threshold, and only one negative NMF component was associated with age, again in females and with an absolute *t* value near 2.

Representative components from PCA and OPNMF showed related developmental anatomy (Figure 3B). PCA component 1 showed decreased expression with age in both males (*t* ≈ − 4.5) and females (*t* ≈ − 6). One pole emphasized lateral/anterior thalamus and precuneus, whereas the opposing pole involved more medial thalamus, occipital cortex, and regions adjacent to the superior central sulcus. Positive OPNMF component 1 showed a similar negative age association (male *t* ≈ − 3; female *t* ≈ − 5), with lateral/anterior thalamus, precuneus, and a more circumscribed field near the lateral sulcus. The similarity in age direction and anatomy suggests that PCA and OPNMF captured a common developmental effect despite representing it with different spatial geometries.

Subject-loading correlations supported this interpretation (Figure 3C). The strongest cross-algorithm correspondences occurred between PCA and OPNMF components, with both positive and negative relationships depending on component orientation. As expected from PCA’s construction, subject loadings from different PCA components were essentially uncorrelated. OPNMF showed a more structured pattern of within-method correspondence, consistent with localized components partitioning a broader source of between-subject variation. Another interpretation is that this could be in part due to the removal of the global scalling, which would capture globally decraesing gray matter. Overall, age sensitivity was concentrated in PCA and OPNMF and differed drastically across algorithms.

### 3.3 Tractography correspondence is highest for positive SC from NMF, OPNMF, and *k*-means

We next compared each cortical SC field with the FA-weighted cortical projection map generated by filtering the thalamocortical tractography atlas through its corresponding thalamic component mask (Figure 4). For PCA and ICA, “positive” denotes same-sign thalamocortical covariance, whereas “negative” denotes opposite-sign covariance. For NMF, OPNMF, and *k*-means, the sign labels identify the positive and negative cortical covariance fields associated with each thalamic component. NMF and OPNMF used a zero-out construction in which values of the opposite sign were set to zero, allowing one thalamic component to be paired with separate positive and negative cortical maps.

**Figure 4.**
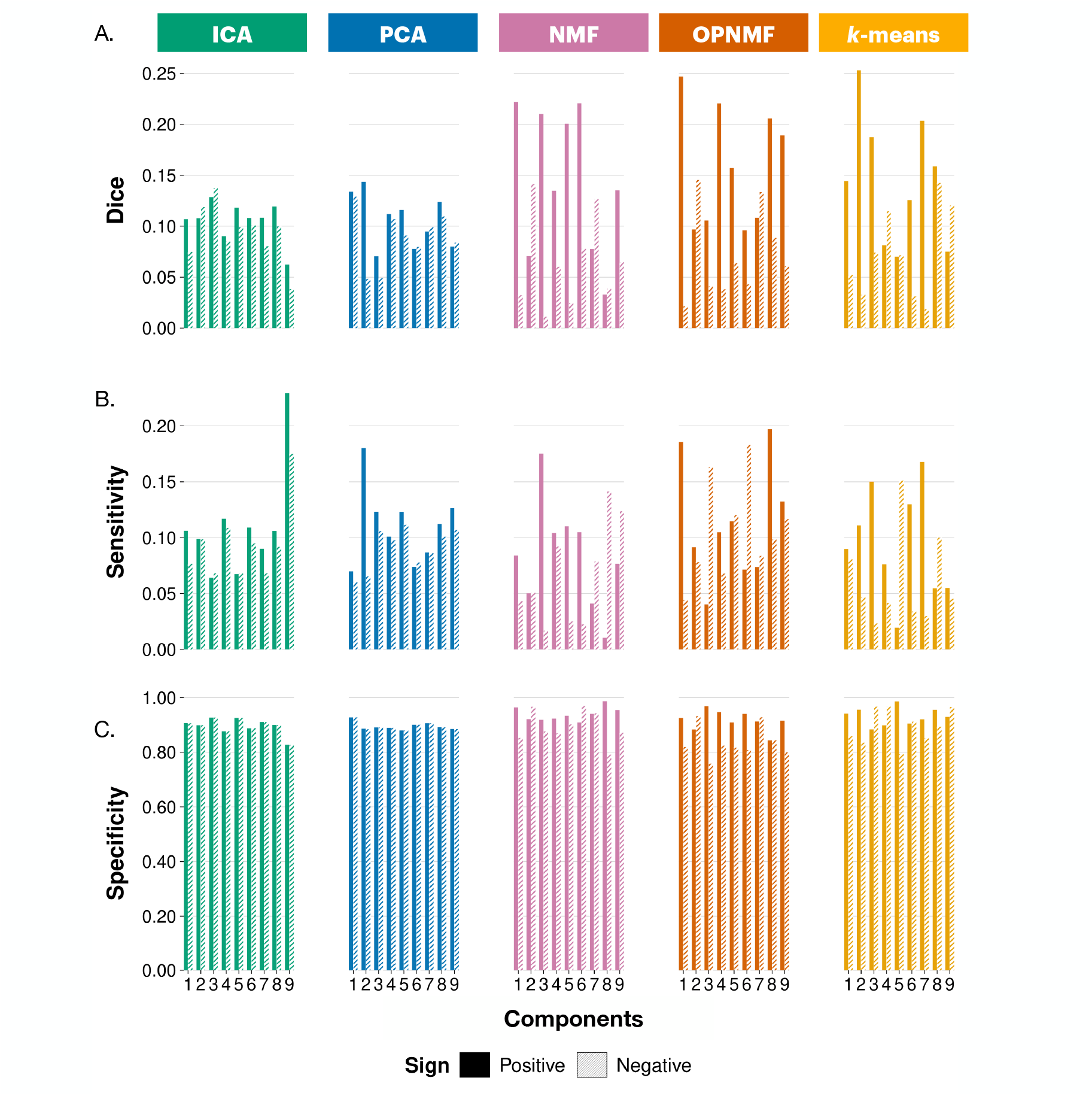
Correspondence between cortical structural covariance fields and tractography-derived thalamocortical projection maps. **A**, soft Dice overlap; **B**, sensitivity; and **C**, specificity. Positive and negative covariance were assessed separately for each method. Soft Dice summarizes graded overlap, sensitivity measures coverage of the tractography-derived field, and specificity measures agreement outside that field.

For spatial overlaps (Figure 4A), a high value of Soft Dice score requires overlap at the right location, high confidence about the magnitudes of values, and high certainty about its background exclusions. NMF, OPNMF, and *k*-means had mean soft Dice values of approximately 0.14–0.16, compared with approximately 0.10 for PCA and ICA. The highest-ranked localized components reached Dice values above 0.20, whereas the best PCA and ICA components remained near 0.14. The ranking reversed for negative covariance: PCA and ICA averaged close to 0.09, while NMF, OPNMF, and *k*-means were lower, generally around 0.06–0.07. Moreover, There is a clear difference of magnitude between positive and negative covariance values throughout, where positive covariance has much higher overlaps in NMF, OPNMF and *k*-means.

Sensitivity was low for every representation and differentiated algorithms less strongly than Dice (Figure 4B). ICA, PCA, and OPNMF means were generally near 0.10–0.12. NMF and *k*-means were lower, approximately 0.05–0.09, with the clearest deficits for negative covariance. Their comparatively strong positive Dice therefore reflects that a restricted subset of the reference-positive cortex was captured by algorithm-derived SC networks.

Specificity revealed a complementary pattern (Figure 4C). For positive covariance, NMF, OPNMF, and *k*-means averaged approximately 0.92–0.94, compared with roughly 0.90 for PCA and ICA. Their positive covariance fields therefore placed less weight outside the tractography-derived territory. Negative covariance again exposed method-specific failure modes. NMF and *k*-means retained low sensitivity, whereas OPNMF specificity fell to approximately 0.83; the other methods remained near 0.89–0.90. This demonstrated that NMF and *k*-means tended to identify small, conservative negative fields, while OPNMF distributed negative covariance across a broader set of tract-negative cortical voxels.

Overall, thalamocortical SC showed only partial correspondence with anatomical connectivity: cortical regions structurally connected to a given thalamic territory were not comprehensively represented in its associated SC field, while covariance also extended beyond the tractography-derived projection territories. For positive covariance, NMF and *k*-means favored compact cortical fields that excluded much of the cortex outside the reference projections, yielding relatively higher Dice and specificity but low sensitivity. OPNMF achieved similarly favorable Dice and specificity while capturing a greater portion of the reference territory, whereas PCA and ICA showed broader cortical coverage at the cost of lower specificity. This trade-off was not preserved for negative covariance, where PCA and ICA showed relatively higher Dice, NMF and *k*-means missed much of the reference territory, and OPNMF exhibited reduced specificity. Together, these findings suggest that SC captures anatomically connected cortical territories selectively rather than comprehensively, with the balance between spatial coverage and localization strongly shaped by algorithmic constraints.

### 3.4 Whole-transcriptome and core–matrix correspondence favour different algorithms

Finally, we examined whether thalamic and cortical territories associated with each SC component exhibited corresponding gene-expression profiles across the transcriptome. For each component, we calculated the component-weighted mean expression of 17,205 AHBA genes separately within the thalamic component and its corresponding cortical SC field. We then assessed the relationship between these two expression profiles across genes, rather than examining spatial correlations between individual genes and SC components. Positive regression slopes indicated that genes with relatively higher expression in the thalamic component also tended to exhibit higher expression in its corresponding cortical field, whereas negative slopes indicated an inverse relationship.

ICA showed the widest range and largest magnitudes of whole-transcriptome regression t-statistics, spanning approximately − 50 to 100 across components (Figure 5B). PCA followed with values ranging from approximately − 20 to 60. The ranges were narrower for OPNMF (approximately − 25 to 40), NMF (approximately − 15 to 30), and *k*-means (approximately − 30 to 20). These results suggest that ICA and PCA identified components with stronger overall thalamocortical correspondence in their weighted gene-expression profiles under this measure. Because the analysis included the entire retained transcriptome, these associations reflect broad transcriptional relationships. These distributions indicate that signed, spatially distributed decompositions preserved stronger broad thalamocortical expression relationships than the localized methods.

**Figure 5.**
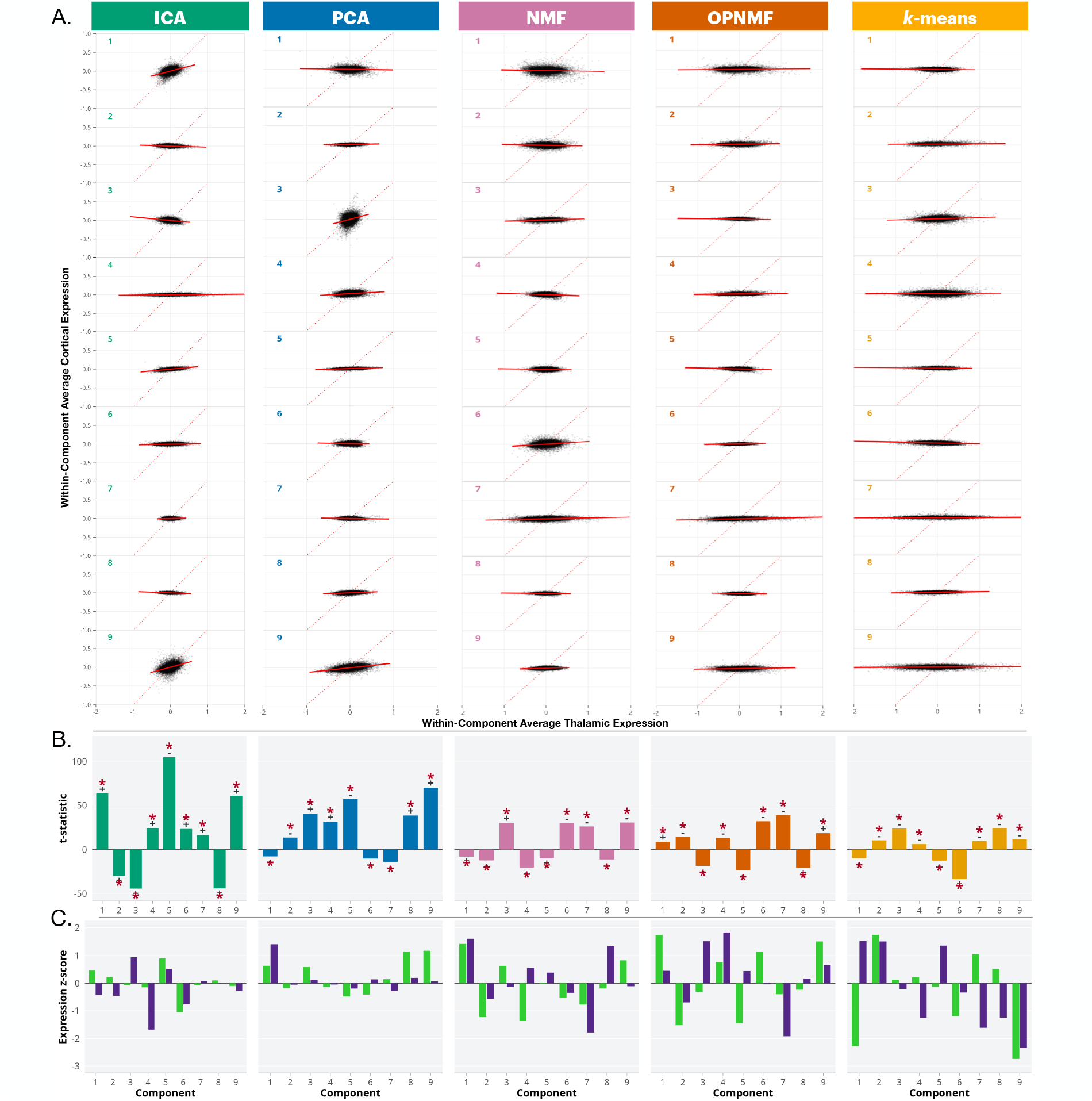
Whole-transcriptome correspondence and targeted core–matrix marker expression. **A**, scatterplots of mean gene expression in each thalamic component against mean expression in its corresponding cortical covariance field, shown for all algorithms, components, and analysed covariance directions. Each point is a retained gene; lines show the fitted linear relationship. Positive slopes indicate concordant thalamic–cortical expression, and negative slopes indicate inverse correspondence. **B**, slope *t* statistics summarizing whole-transcriptome correspondence. Where a single value is displayed per component, the covariance direction with the larger absolute statistic is retained and its sign is preserved; this dominant-direction summary is descriptive because direction is selected from the observed data. **C**, standardized thalamic expression of *PVALB* and *CALB1* for the dominant covariance direction of each component. Scores are *z*-standardized separately by gene across the full set of algorithm–component combinations; positive values therefore indicate relatively high marker expression within this comparison set, not significance against a spatial null.

The targeted marker analysis showed a different pattern (Figure 5C). Here, we examined the component-weighted mean thalamic expression of *PVALB* and *CALB1*, with each marker standardized separately across all algorithm-component combinations. NMF, OPNMF, and *k*-means showed relatively higher marker expression across more components, whereas PCA and ICA tended to have lower scores. This indicates that the thalamic regions emphasized by the localized methods tended to have higher average expression of one or both core-matrix-associated markers. Here, lower scores reflect either consistently low marker expression or averaging across territories with different expression levels.

The two transcriptomic analyses therefore capture different properties of the components. The whole-transcriptome analysis examined the correspondence between thalamic and cortical expression profiles across thousands of genes, whereas the targeted analysis measured the relative expression of two selected markers within the thalamic component maps. Their different rankings may inform how the algorithms distribute component weights across molecularly distinct territories, without necessarily implying that localized methods recover more homogeneous cell populations.

## 4 Discussion

In this work, we compared five commonly used data decomposition algorithms on DBM-derived thalamocortical SC matrices to study how the resulted SC networks differ systematically in their ability to capture different aspects of biology. At the phenotypic level, we identified PCA and OPNMF as optimal choices for the concentration of age-related variances. At the connectivity level, we found superior performances in methods with additive and partition-based representations, i.e., NMF, OPNMF, and *k*-means. At the transcriptomics level, we showed diverging results for whole-transcriptomics and marker genes that are dependent on the spatial representations of each algorithm.

### 4.1 Algorithmic geometry selects biological scales

First, our findings support that each method selects a unique combination of biological information. PCA and OPNMF were most sensitive to age-related variation. NMF, OPNMF, and *k*-means most closely matched positive tractography-derived cortical territories. ICA and PCA showed the broadest whole-transcriptome correspondence, while NMF, OPNMF, and *k*-means localized relatively *PVALB*- and *CALB1* -rich thalamic fields. These divergent results rule out a simple hierarchy from weak to strong methods. The algorithms exposed complementary views of the same population covariance because each rewarded different spatial and statistical properties.

This points to a nuanced interpretation of SC networks through the interplay between biological patterns and algorithmic optimization rules in dimensionality reduction. PCA rewards covariance that explains population variance along orthogonal axes; ICA rewards separable non-Gaussian sources; NMF rewards additive non-negative parts; OPNMF rewards a projective and competitive version of those parts, and *k*-means rewards mutually exclusive territories. Biological benchmarks are also geometrically distinct: developmental effects can be broad and smoothly varying; tractography-derived projection maps are localized and modular; whole-transcriptome organization contains broad, spatially correlated gradients; and individual markers can be concentrated in circumscribed fields. Consequently, evaluation performance increases when the geometry of the method resembles the geometry of the biological benchmark. Algorithmic constraints therefore implicitly operate as priors on biological scales.

This interaction is particularly clear in thalamocortical SC, where biological organization is neither purely modular nor purely continuous. Thalamic nuclei and their preferential cortical projections support a modular description, whereas transcriptomic and connectomic gradients can extend across nuclear boundaries (Oldham and Ball, 2023; John et al., 2025). Consistent with this mixed organization, all methods recovered major, recognizable thalamocortical systems, but differed in how sharply they partitioned those systems and which within-system variations they preserved. These differences reflect distinct representational emphases that may be informative for different biological questions. The relevant comparison is therefore not which algorithm provides a more faithful account of thalamocortical organization, but which aspects of that complex organization each method rewards and how they relate to the biological benchmark under study.

### 4.2 Orthogonality combined with variance optimization captures broad developmental variation

Developmental SC has been interpreted partly as coordinated maturation across distributed systems, and human thalamocortical connectivity continues to reorganize through adolescence along a sensorimotor-to-association hierarchy (Alexander-Bloch et al., 2013b; Sydnor et al., 2025). The age analysis here suggests that dominant neurodevelopmental variation in this sample was spatially broad enough to be captured efficiently by PCA and, possibly through a different mechanism, OPNMF.

In PCA, a signal that changes consistently across many voxels contributes strongly to a leading PCA axis even when no single territory carries it. PCA’s sensitivity to age is therefore compatible with its variance-maximizing objective.

OPNMF does not maximize variance in the same way, yet it may concentrate the same broad source by limiting redundant spatial representation. Its non-negative projective basis yields localized maps, while orthonormality forces components to compete in variance optimization. A developmental pattern distributed across the thalamus and cortex may consequently be divided into a small set of non-redundant territories whose subject scores remain strongly age-sensitive. Large population studies have shown that projective non-negative components can be stable across model scales and useful for biological association analyses (Wen et al., 2023). Here, the strong correlation between PCA and OPNMF subject loadings, together with similar representative anatomy, suggests that the two methods encoded similar developmental information despite their different spatial forms.

Orthogonality is the key shared factor between PCA and OPNMF, yet the result does not imply that orthogonality alone identifies development, since it was also present in the whitening step of ICA, which showed very few age associations. A methodological interpretation is that global, smooth developmental variation was captured by the low redundancy from orthogonality constraints in PCA and OPNMF combined with their particular optimization scheme.

Additionally, the age-sensitive SC networks, along with their sex-stratified concordance could reveal some generalizable knowledge about thalamocortical structural coupling during adolescence to young adults. Take PC1 and positive OPNMF component 1 as examples: PC1 crossed from positive to negative loadings with age, marking a population-level shift from a lateral-anterior thalamus–precuneus configuration toward an opposing medial thalamus–occipital-pericentral configuration. OPNMF component 1 remained positive but declined in expression, isolating a decreasing contribution of the former configuration as the age increases. Their convergence may reflect adolescent rebalancing among thalamocortical systems with different maturational timing embedded in morphological concordance. Components detected in both females and males indicate that the principal age directions were not obviously driven by one sex.

### 4.3 Additive and partition-based representations recover compact positive projection fields

The connectivity benchmark favoured a different set of constraints. What NMF, OPNMF, and *k*-means share is a tendency to limit spatial mixing. Conceptually, NMF builds additive parts, OPNMF makes those parts compete under a projective orthonormal representation, and *k*-means assigns each voxel to one territory. Structural connectivity from tractography has a compatible geometry: after filtering through a thalamic mask, cortical projection weights are spatially bounded. A localized component can therefore overlap a connectivity-based projection field without carrying the broad opposing lobe that is natural to a signed PCA or ICA axis. This spatial precision is also consistent with the parts-based interpretation originally motivating the use of NMF in structural neuroimaging (Sotiras et al., 2015).

Dice, sensitivity, and specificity should be interpreted jointly. Here, sensitivity prevents an overly generous reading. NMF and *k*-means covered only a small fraction of the tractography-derived cortical field. Their relatively high Dice and specificity are best understood as identification of a conservative projection core with few false-positive voxels. OPNMF offered a somewhat broader coverage for positive covariance, but its negative specificity was poor. Because most cortical voxels lie outside any one thalamic projection map, specificity can also look high under substantial mismatch.

The pattern of overall worse performance in negative covariance may reveal how these constraints handle anticovariance. To elaborate, NMF and *k*-means had low negative sensitivity, consistent with a diffuse or weak anticovariance field being difficult to express as additive parts or a hard cluster-associated map. OPNMF retained sensitivity but lost specificity, suggesting that projective competition may redistribute negative covariance over a broad cortical field when components are forced to remain non-negative and comparatively orthogonal.

### 4.4 Distributed methods preserve broad transcriptomic structure; localized methods isolate selected core-matrix markers

Transcriptomic results revealed a scale-dependent contrast between algorithms. ICA and PCA produced the largest-magnitude whole-transcriptome statistics, whereas NMF, OPNMF, and *k*-means produced higher relative *PVALB* and *CALB1* expression. The adult human brain transcriptome is strongly spatially structured (Hawrylycz et al., 2012), and thalamic expression varies along axes that cross nuclear boundaries and correspond to cortical connectivity (Oldham and Ball, 2023). PCA can preserve dominant shared gradients, while ICA can separate distributed sources that overlap anatomically but differ statistically. Their stronger *t* statistics may therefore reflect sensitivity to broad transcriptional relationships rather than precise localization of a single cell population.

The marker analysis rewards almost the opposite property. Localized NMF, OPNMF, and *k*-means components reduce averaging between marker-rich and marker-poor tissue. Higher *PVALB* and *CALB1 z* scores may thus reflect sharper localization of core- and matrix-associated fields. This interpretation is consistent with with work relating the relative expression of these calcium-binding proteins to thalamocortical functional gradients and cortical timescales (Müller et al., 2020), as well as multimodal evidence that a secondary thalamocortical connectivity axis follows core–matrix-associated variation (John et al., 2025).

The two transcriptomic analyses are complementary. Whole-transcriptome correspondence asks whether a component preserves a relationship spread across many genes; the marker analysis asks whether it isolates selected, molecularly differentiated territories. A method may perform well on one while deliberately smoothing or fragmenting the other. More generally, “transcriptomic correspondence” is not a unitary property. A broad transcriptional gradient, a co-expression module, a cell-type signature, and a single marker each impose different spatial targets.

### 4.5 Recommendations on aligning algorithms with the biological questions

Overall, the five algorithms tested showed clear trade-offs across biological aspects and scales, with no single method optimal for all objectives. PCA is a strong candidate when the target is dominant developmental variation or a broad transcriptomic gradient. ICA is preferable when the biological signal is expected to be distributed and statistically separable. NMF is useful for compact positive projection fields and focal genetic marker-rich territories, with the cost of low coverage. OPNMF excels in age sensitivity, positive tract overlap, and marker localization, but its negative specificity shows that it is not uniformly robust. *k*-means provides the clearest territories and strong positive anatomical/marker correspondence, but sacrifices overlap and broad subject-level variation.

When the target geometry is uncertain, explorative analyses with a few different algorithms is recommended: apply at least one distributed signed method and one localized additive or partition-based method. Agreement across them is less representation-dependent, while disagreement can reveal whether a phenomenon behaves more like a gradient, overlapping system, additive part, or discrete territory. Naming components by their derivation, e.g., “OPNMF-derived SC”, would make this dependence explicit.

In practice, we also recommend testing and visualizing a few numbers of components that have reasonably good metrics, and determine the number of components with judgment considering the goal of the study. Similarly, exploring different decomposition parameter choices might be helpful for understanding the structure of the data.

### 4.6 Conclusion

Broadly, our results emphasize that algorithm selection in SC studies is a representational choice with inherently biological consequences. We suggest future work to align decomposition choices with explicit biological targets and avoid cross-study comparisons that ignore algorithmic differences. By mapping these mathematical constraints onto biological readouts, this work motivates SC studies toward a more principled and question-driven use of decomposition algorithms. Although our work focuses on thalamocortical SC and the precise balance of biological influences may differ for other structures, the general algorithm-specific profiles observed in this study are likely to extend in other SC networks. We hope this work can motivate a broader shift toward explicitly studying voxelwise subcortical-cortical SC networks, and towards interpreting their biological relevance in light of the specific choice of algorithm used upstream to the analyses.

### 4.7 Limitations

Overall, our results show that biological correspondence of SC depends on the implemented representation and reference biology in the thalamocortical system. These findings motivate explicit alignment between methodological choices, biological questions, and the spatial and statistical properties of the benchmarks used to evaluate them. The limitations of this study are related to methodological decisions, innate comparability between algorithms, and the characteristics from the resources used for biological references warrant consideration.

#### 4.7.1 Methodological decisions

The present comparison is circumscribed by the set of choices concerning algorithmic parameters, number of components and clusters, and matrix construction. Note that our study does not provide an exhaustive assessment of each method’s best achievable performance for any dataset. In particular, algorithm implementation parameters like initialization, regularization, whitening, and convergence criteria can affect the solutions obtained. For example, explicit sparsity constraints or penalties can alter the localization and overlap of NMF components (Hoyer, 2004).

The consensus number of components, selected by considering reconstruction metrics and anatomical organization, provides a common reporting scale but may not be optimal for every method. Our anatomical appraisal introduces an additional interpretive choice, potentially favoring recognizable territories. PCA’s variance ordering also distinguishes the interpretation of leading components from assessment of the full retained representation (Jolliffe and Cadima, 2016). Consequently, our conclusions about individual components or method rankings remain conditional on the selected model order.

Matrix construction also affects the biological variation available to the algorithms. Relative log-Jacobians emphasize local morphology, while subsequent structure-volume residualization conditions the measurements on thalamic or cortical size. Global-effect removal can also change both covariance patterns and the occurrence of negative correlations (Zhang et al., 2025). Furthermore, we have observed that upstream decisions concerning quality control, intensity correction, registration, and masking introduce could also affect the results to some extent and SC estimates are demonstrated to be sensitive to acquisition and processing choices (Carmon et al., 2020). In particular, residual misalignment, partial-volume effects, and smoothing may blur regional boundaries or introduce shared measurement variation. Although we did not investigate the exact consequences of each processing choice, we recommend considering processing choices carefully prior to analyses.

While recognizing the potential differences in results from alternative preprocessing, processing schemes and even using different morphometric measures, at the broad scale of our analyses, our main suggestion about the alignment between algorithmic and biological representations remain generally adaptable.

#### 4.7.2 Comparability across algorithms

It is important to highlight that the algorithms tested do not produce interchangeable spatial quantities, thus a perfectly fair comparison is nearly impossible to achieve. Our attempt to make them comparable by normalizing maps to a common numerical range facilitates spatial comparison, but does not address the innate differences in their representations of SC networks. PCA and ICA generate signed component contributions whose global orientation is arbitrary; when the input is centered, these contributions describe deviations from the mean covariance profile rather than necessarily the sign of the original correlations. NMF and OPNMF instead represent positive and negative-magnitude input channels additively. Thalamic *k*-means assignments encode binary membership, whereas their cortical centroids are continuous signed averages. Moreover, separating signs before averaging or factorization is not equivalent to separating the signs of an already averaged profile. Thus, the terms “positive-” and “negative SC” do not signify identical statistical meanings across methods.

#### 4.7.3 Alignment of study data and resources

It is worth highlighting that the anatomical, connectivity, and transcriptomic resources provide complementary references rather than ground truth for the biological processes generating SC.

Although the study-specific PNC template reduces dependence on a fixed adult anatomical target, mapping external atlases to this template cannot eliminate differences in anatomy, age, resolution, or modality. Registration and resampling may be especially consequential near cortical boundaries and small thalamic nuclei.

The thalamocortical tractography atlas was constructed from an HCP Young Adult group-average diffusion template; developmental datasets, including PNC and HCP-D, were subsequently used to evaluate its reconstruction in individuals, and reconstructability in these datasets also informed atlas curation (Sydnor et al., 2025). It is an external anatomical resource but not entirely independent of the broader PNC data resource. The FA reference has an additional spatial-resolution limitation. Since each HCP-MMP cortical parcel was assigned the mean FA of its atlas-defined thalamic connection, giving the same microstructural weight to all voxels within that parcel, our voxelwise comparison can remain influenced by parcel boundaries and connection-level averaging. Accordingly, our findings here should be interpreted as correspondence with this atlas, weighting scheme, and seed definition.

As for the AHBA, its microarray resource comprises six adult donors aged 24–57 years, with unequal hemispheric coverage and a strong male predominance (Hawrylycz et al., 2012; Markello et al., 2021). These characteristics warrant consideration for the transcriptomics analysis given the age and sex mismatch with the sample population, since adolescent brains might express a slightly different transcriptomic landscape. For example, prefrontal microarray analyses have reported distinct late-adolescent expression profiles involving *MBP, MPZL1*, and the axon-guidance gene *PLXND1* (Harris et al., 2009). Transcriptomic correspondence also depends on probe selection, tissue sampling, spatial assignment, normalization, and donor aggregation (Arnatkevičiūtė et al., 2019; Markello et al., 2021).

## Author contributions

Y.L., M.M.C., and Y.Y. conceptualized the study. Y.L., V.J.S., M.M.C., and Y.Y. developed the methodology. Y.L. developed the analysis software, curated the data, performed the analyses and validation, prepared the visualizations, and coordinated the project. Y.Y. contributed to the analyses. A.D. preprocessed the Philadelphia Neurodevelopmental Cohort data and provided participant ages in decimal years. J.Z. and S.F. contributed to deformation-based morphometry data quality control. G.A.D. provided methodological and technical support for deformation-based morphometry. V.J.S. contributed to the connectivity analyses. Y.L. drafted the manuscript. Y.L., V.J.S., A.D., T.D.S., M.M.C., and Y.Y. critically revised the manuscript. All authors reviewed and approved the final version of the manuscript.

## Acknowledgements

The authors declare no competing interests.

The authors acknowledge the contributions of Manuela Costantino and Swapna Premasiri to preprocessing data quality control.

