## Supplementary Methods for "Algorithmic Geometry Shapes the Biological Meaning of Thalamocortical Structural Covariance"

Algorithmic representations of thalamocortical structural covariance

#### S1 Acquisition, image quality control, and analytical cohort

##### S1.1 MRI acquisition and phenotype data

T1-weighted (T1w) MRI and phenotype data were obtained from the Philadelphia Neurodevelopmental Cohort (PNC) (Satterthwaite et al., 2014). The structural release contained 1,598 images from participants aged 8–22 years. Images were acquired at the Hospital of the University of Pennsylvania on a Siemens 3 T TIM Trio scanner with a 32-channel head coil. The magnetization-prepared rapid acquisition gradient-echo sequence used a repetition time of 1,810 ms, echo time of 3.51 ms, inversion time of 1,100 ms, flip angle of 9°, and approximate voxel dimensions of $0.94 \times 0.94 \times 1.00\,\text{mm}^3$. Age, sex, and medical-history variables were taken from the curated PNC phenotype tables.

##### S1.2 Motion and intensity-processing quality control

All raw T1w images were visually inspected for motion-related artifacts using the laboratory motion-quality-control protocol[1]. Images receiving an ordinal rating of 2 or less were retained, yielding 1,097 images.

These images were processed with the iterativeN4 pipeline[2], which combines N4 intensity non-uniformity correction with iterative tissue classification (Tustison et al., 2010). Outputs comprised a bias-corrected T1w image, tissue classification, and brain mask. Outputs were visually assessed for brain coverage and gross processing failure; the first 100 cases received an additional independent review. All 1,097 images passed this stage and entered population-template construction.

##### S1.3 Region-focused registration quality control

After registration, each participant's resampled T1w image was compared with the PNC population template within the atlas-defined thalamic and cortical regions. Images were excluded for gross misregistration that substantially displaced either structure relative to the template. This was an analysis-specific assessment of the thalamus and cortex, not comprehensive whole-brain registration quality control. A total of 1,028 images passed. Restriction to medical-history ratings of 0 or 1 and availability of the phenotype variables required for analysis produced a final cohort of 755 participants.

[1] https://github.com/CoBrALab/documentation/wiki/Motion-Quality-Control-(QC)-Manual

[2] https://github.com/CoBrALab/iterativeN4_multispectral

Table S1: Sample-retention sequence.

| Stage | Images/participants retained |
|---|---|
| PNC structural MRI release | 1,598 |
| Raw-image motion quality control | 1,097 |
| Intensity-processing quality control and template-building input | 1,097 |
| Thalamic/cortical registration quality control | 1,028 |
| Medical-history and phenotype eligibility | 755 |

### S2 DBM template construction and spatial references

#### S2.1 Population-template construction and Jacobian maps

Deformation-based morphometry (DBM) relates local volumetric differences to the Jacobian determinants of nonlinear registration transformations (Chung et al., 2001). Bias-corrected images were processed using the single-level workflow implemented in `optimized_antsMultivariateTemplateConstruction` (Germann et al., 2025). Initialization used the ICBM152 nonlinear symmetric 2009c T1w template and corresponding brain mask; participant brain masks guided registration.

The PNC population template was generated over six iterations. At each iteration, participant images underwent affine and symmetric diffeomorphic nonlinear registration (SyN) to the current target in ANTs (Avants et al., 2008, 2011). Template intensities were updated from the registered images, and template shape was updated using their average affine and nonlinear transformations. The pipeline's intensity-sharpening operation occurred within this iterative construction, rather than as a separate modification of the completed population reference. The updated template served as the next registration target.

Relative Jacobian determinant maps were generated from the final template-building iteration. These exclude the affine volume contribution and remove residual affine effects within the nonlinear deformation field, in contrast to absolute Jacobians that retain affine and nonlinear contributions (Germann et al., 2025). The relative maps were log-transformed. Under the pipeline convention, positive and negative log-Jacobian values indicate locally larger and smaller anatomy, respectively, relative to the population reference after affine adjustment. Separate outputs were generated with Gaussian smoothing full widths at half maximum of 1, 2, and 4 mm.

#### S2.2 Allen reference-atlas registration

The Allen Human Reference Atlas–3D (AHRA) label volume is defined on the ICBM152 nonlinear symmetric 2009b anatomical reference (Ding et al., 2016, 2020). This is distinct from the 2009c image used to initialize the PNC template. In ANTs version 2.5.4, the PNC population template was the fixed image and the AHRA-associated 2009b image was the moving image. Registration was initialized from relative image centres and proceeded through rigid, affine, and nonlinear SyN stages. The rigid and affine stages used mutual information with 32 histogram bins and regular sampling of 25% of voxels. The nonlinear stage used cross-correlation with a radius of four voxels.

The resulting affine and nonlinear transforms were applied to the AHRA annotation volume using `antsApplyTransforms`, with the PNC template as the reference image and MultiLabel interpolation for discrete labels. The transformed anatomical reference and annotations were overlaid on the

PNC template for visual inspection at the available resolutions.

##### S2.3 Masks and spatial sampling

AHRA ontology entries belonging to the thalamus and cerebral cortex were combined across hemispheres to create bilateral masks. The voxel counts were:

Table S2: Anatomical masks and voxel counts.

| Resolution | Thalamic voxels | Cortical voxels |
|---|---|---|
| Higher resolution | 10,707 | 661,187 |
| Lower resolution | 1,322 (not used) | 82,603 |

#### S3 Structure-volume adjustment and matrix definitions

##### S3.1 Residualization

Let $J_{it}^{T}$ and $J_{ic}^{C}$ denote the log-transformed relative Jacobian values for participant $i$ at thalamic voxel $t$ and cortical voxel $c$. Voxelwise linear models were fitted across participants:

$$J_{it}^{T} = \beta_{0t} + \beta_{1t} V_i^{T} + \epsilon_{it}^{T}, \tag{S1}$$

$$J_{ic}^{C} = \beta_{0c} + \beta_{1c} V_i^{C} + \epsilon_{ic}^{C}, \tag{S2}$$

where $V_i^T$ and $V_i^C$ are the corresponding total structure volumes. Unstandardized residuals were retained as the adjusted measurements. This adjustment was performed separately for the thalamus and cortex, in addition to excluding affine effects when generating relative Jacobians.

##### S3.2 Group-level SC matrix

With $N = 755$, adjusted data were arranged into participant-by-voxel matrices

$$\mathbf{X}_T \in \mathbb{R}^{N\times V_T}, \qquad \mathbf{X}_C \in \mathbb{R}^{N\times V_C}. \tag{S3}$$

Pearson correlations across participants defined the thalamic-by-cortical SC matrix:

$$R_{tc} = \mathrm{corr}(\mathbf{X}_T[:,t], \mathbf{X}_C[:,c]), \qquad \mathbf{R} \in [-1,1]^{V_T\times V_C}. \tag{S4}$$

Each row was a thalamic voxel's cortical SC profile. Positive and negative entries indicated concordant and opposing structure-volume-adjusted morphological differences across participants. Entries were correlations, rather than unstandardized covariance estimates or participant-specific connections.

##### S3.3 Participant-level input

A separate joint morphometric matrix preserved individual measurements:

$$\mathbf{Y} = [\mathbf{X}_T \mid \mathbf{X}_C^{(d)}] \in \mathbb{R}^{N\times(V_T+V_C^{(d)})}, \tag{S5}$$

where $\mathbf{X}_C^{(d)}$ contains spatially downsampled cortical features. In the spatial-row convention,

$$\mathbf{Y}^\top \in \mathbb{R}^{(V_T+V_C^{(d)})\times N}. \tag{S6}$$

Spatial factors describe joint thalamic/cortical voxel patterns, and the complementary factors index participants. These are separate fits from the decompositions of $\mathbf{R}$.

##### S3.4 Non-negative encodings

For the spatial SC fits, positive and negative-magnitude cortical features were concatenated:

$$\mathbf{R}^+ = \max(\mathbf{R}, 0), \qquad \mathbf{R}^- = \max(-\mathbf{R}, 0), \tag{S7}$$

$$\mathbf{R}_\pm = [\mathbf{R}^+ \mid \mathbf{R}^-] \in \mathbb{R}_+^{V_T\times 2V_C}. \tag{S8}$$

NMF and OPNMF were fitted jointly to this non-negative encoding. Their cortical factors were subsequently divided into positive- and negative-covariance blocks. For participant-level analyses, the two signs were instead fitted independently:

$$\mathbf{Y}^+ = \max(\mathbf{Y}, 0), \qquad \mathbf{Y}^- = \max(-\mathbf{Y}, 0). \tag{S9}$$

The two matrices were oriented with spatial observations for their separate fits. These signs identify adjusted morphometric values, not positive and negative correlations. With $K = 9$ in each fit, each non-negative method provides two separately estimated participant-level sets, not one nine-component representation identical in flexibility to signed PCA or ICA.

**Constraint scope.** In the thalamus-row SC formulation, sign splitting doubles the cortical columns of $\mathbf{R}$; it does not create positive/negative thalamic rows. If the OPNMF spatial basis is $\mathbf{W} \in \mathbb{R}_+^{V_T\times K}$, its orthogonality constraint acts on those thalamic weights. Under exact non-negative orthogonality, a row cannot have positive entries in two basis columns because their non-negative inner product would then be positive. Numerical fits may approximate this property.

#### S4 Algorithm settings and model-order selection

##### S4.1 Software and numerical implementation

PCA, ICA, NMF, and $k$-means used scikit-learn version 1.7.1 (Pedregosa et al., 2011). OPNMF used `juaml/opnmf`[3]. Objective functions are described in the main Methods. Implementation settings are detailed below.

**PCA.** A randomized solver was used to accommodate matrix size (Jolliffe and Cadima, 2016).

**ICA.** Cortical features were centred across thalamic voxels before custom whitening. Randomized singular value decomposition retained $K$ dimensions with 15 oversamples, four power iterations, and random state 0. FastICA used the parallel algorithm, `logcosh` contrast, a maximum of 5,000 iterations, tolerance $10^{-4}$, and random state 0. Internal FastICA whitening was disabled because whitening had already been performed. The ICA stage used single-precision values, and cortical coefficients were mapped from whitened coordinates back to the original cortical-feature space (Hyvärinen and Oja, 1997, 2000).

[3] `https://github.com/juaml/opnmf`

**NMF.** The sign-split matrix was fitted using coordinate descent with NNDSVD initialization, a maximum of 5,000 iterations, tolerance $10^{-4}$, and random state 0 (Lee and Seung, 1999).

**OPNMF.** The sign-split matrix was fitted using NNDSVD initialization, a maximum of 50,000 iterations, and tolerance $10^{-5}$ (Yang and Oja, 2010; Sotiras et al., 2015).

***K*-means.** Signed cortical profiles were clustered using Lloyd's algorithm, $k$-means++ initialization, `n_init="auto"`, a maximum of 10,000 iterations, tolerance $10^{-4}$, and random state 0. Input values were single precision (Lloyd, 1982; Arthur and Vassilvitskii, 2007).

##### S4.2 Selection of component numbers

The candidate range was $K = 2$–25. Reconstruction diagnostics were evaluated for PCA, ICA, NMF, and OPNMF using a fixed 10% entry-level hold-out mask, with scores taken from the tenth matrix-completion iteration. Clustering diagnostics were evaluated for $k$-means. A common model order of $K = 9$ was used for the cross-algorithm comparison (Figure S1). This was retained as a common reporting scale, not a demonstrated optimum for every algorithm.

#### S5 Anatomical characterization and map-overlap definitions

##### S5.1 Atlas labels and map normalization

AHRA labels were selected using ontology paths. Thalamic labels came from the diencephalon/thalamus branch; cortical labels came from the cerebral cortex and cerebral gyri and lobules branches. Individual labels were retained and intersected with the corresponding high-resolution mask rather than merged into larger regions.

For a signed spatial map $\mathbf{u}$, the two fields were obtained as

$$u_v^+ = \max(u_v, 0), \qquad u_v^- = \max(-u_v, 0). \tag{S10}$$

Each nonempty field was divided by its own maximum; empty fields remained zero. NMF/OPNMF thalamic maps and their separate cortical channels were maximum-normalized in the same way. Thalamic $k$-means memberships were binary; signed cortical centroids were separated into positive and negative-magnitude fields. OPNMF winner-take-all maps were for display only, with continuous weights retained for quantitative analyses.

##### S5.2 Atlas overlap

Spatial overlap between normalized component maps and binary AHRA label masks was quantified using soft Dice scores.

##### S5.3 Heatmaps and representative maps

Positive/negative labels identified a component side, not a negative overlap score. For each component–atlas-label pair, the higher of the two side-specific overlap scores was shown, with ties assigned to the positive side. Selection occurred separately for each atlas label, so a heatmap row

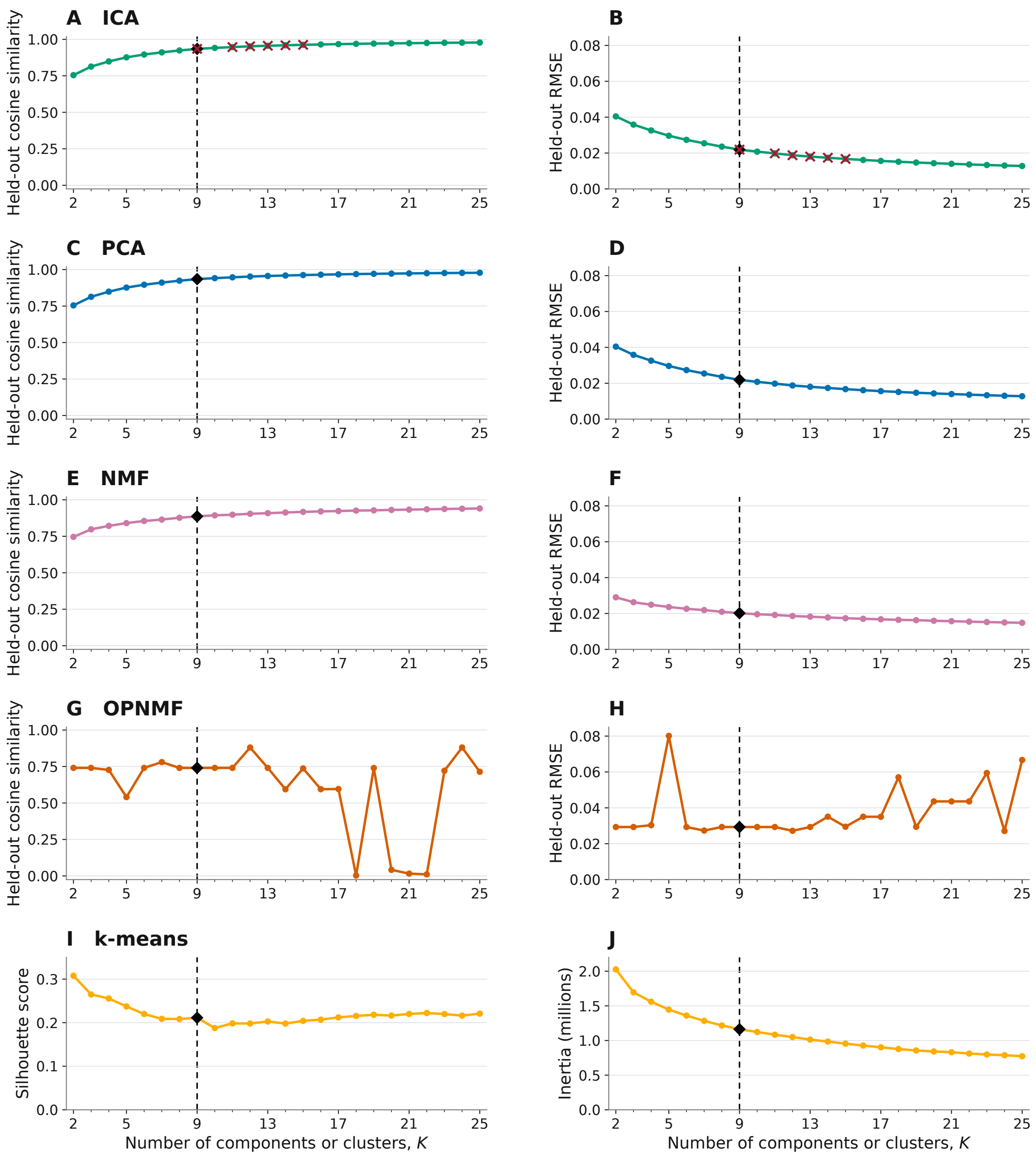


Figure S1: **Reconstruction and clustering diagnostics across component counts.** Held-out cosine similarity (left) and root mean squared error (RMSE; right) are shown for ICA (A–B), PCA (C–D), NMF (E–F), and OPNMF (G–H). For $k$-means, silhouette score (I) and within-cluster inertia (J) are shown. For the factorization methods, each point represents the tenth iteration of matrix completion using a fixed 10% entry-level hold-out mask. These iterations are successive updates rather than independent hold-out replicates; error bars are therefore not shown. Dashed lines and black diamonds mark $K = 9$, retained as the common reporting scale for cross-algorithm comparisons rather than a demonstrated optimum for every method. Red crosses indicate final ICA fits flagged as non-converged. Higher cosine similarity and silhouette scores, and lower RMSE and inertia, indicate better performance for the respective metric.

need not describe a single global component pole. Atlas-label ordering was fixed across algorithms within each structure. Cortical labels were grouped according to ontology for display only, without pooling masks or scores. Allocortex and periallocortex display groups were omitted from the main heatmap but retained in the overlap tables. Positive or unsigned/binary maps were red and negative-side maps were blue, with intensity indicating overlap. Color limits were shared across algorithms within each structure, with separate limits for thalamus and cortex.

Representative maps were selected by spatial Pearson correspondence with PCA component 1.

### S6 Participant-weight associations with age

#### S6.1 Weights and regression models

Participant weights were obtained from the separate fits to the joint morphometric input. PCA/ICA weights were signed. NMF/OPNMF weights were non-negative and analysed separately for positive- and negative-value fits. Several $k$-means solutions in this pathway were confined to one anatomical structure; this implementation was excluded from the primary paired thalamocortical participant-weight comparison. This exclusion does not imply that participant summaries can never be constructed from $k$-means clusters.

Within each sex, a separate model was fitted for each algorithm/component:

$$L_{ik} = \beta_{0k} + \beta_{1k}\,\mathrm{Age}_i + \epsilon_{ik}, \tag{S11}$$

where $L_{ik}$ is the participant's component weight. The signed age-coefficient statistic was

$$t_k = \frac{\widehat{\beta}_{1k}}{\mathrm{SE}(\widehat{\beta}_{1k})}. \tag{S12}$$

The sign of a PCA/ICA association was interpreted relative to the retained spatial orientation. For NMF/OPNMF, the two input-sign solutions remained separate. Pairwise correlations of participant-weight vectors within and across algorithms assessed shared individual differences.

#### S6.2 Interpretation of age associations

These are cross-sectional age associations, not estimates of within-person change or out-of-sample age-prediction performance. The $t$ statistic combines association magnitude and uncertainty; it is not an effect-size estimate independent of sample size. A significant association in one sex but not the other does not constitute a test of different slopes.

### S7 Tractography reference construction and spatial comparisons

#### S7.1 Spatial transformation, seed masks, and filtering

The connectivity reference was the PennLINC thalamocortical tractography atlas in HCP1065 1.25-mm space (Sydnor et al., 2025). Algorithm-derived thalamic and cortical maps were transformed from the PNC template into this reference space.

For PCA, ICA, NMF, and OPNMF, the highest-weight 15% of thalamic voxels were retained as seed masks. PCA/ICA used upper- and lower-tail masks to represent both poles. This threshold was chosen to make continuous-map seed selection comparable while limiting the fragmentation seen at more restrictive thresholds. Binary $k$-means clusters were not percentile-thresholded and thus retained their native, potentially different sizes.

Each mask filtered atlas streamlines in DSI Studio version `chen-2023-02-27` (Yeh et al., 2013). In the region-of-interest condition, a streamline was retained if it intersected the mask. In the endpoint condition, a streamline was retained if an endpoint lay within the mask. Filtered streamlines were exported as cortical tract-density images (TDI).

##### S7.2 Signed map comparisons

Cortical SC maps were separated into positive and negative-magnitude fields and normalized to $[0, 1]$ within a common cortical mask. PCA/ICA same-sign thalamic and cortical poles defined positive component contributions and opposite-sign poles defined negative contributions. For NMF/OPNMF, each thalamic component was paired with its separate positive and negative cortical channels. For $k$-means, the cortical centroid supplied the two signed fields.

Agreement between continuous map weights describes weighted spatial correspondence; it does not count anatomically true or false connections. Normalization to $[0, 1]$ does not make the weights calibrated probabilities.

#### S8 AHBA processing and transcriptomic summaries

##### S8.1 Expression resource and spatial assignment

The Allen Human Brain Atlas (AHBA) supplied post-mortem expression measurements (Hawrylycz et al., 2012). Thalamic and cortical tissue samples were processed and normalized separately. Probe filtering and annotation retained 17,205 genes. The refined sample localization described by Yee et al. (2025) was used to relate AHBA expression measurements to the PNC population-template space.

##### S8.2 Component-weighted expression

Component-weighted mean expression was calculated for each retained gene in the thalamus and cortex. Continuous spatial weights were used for factorization methods and binary membership for thalamic $k$-means clusters. The cortical profile of a $k$-means cluster is signed and continuous.

Let $E^T_{gakd}$ and $E^C_{gakd}$ denote the component-weighted mean expression of gene $g$ in the thalamus and cortex, respectively, for algorithm $a$, component $k$, and covariance direction $d$.

##### S8.3 Whole-transcriptome correspondence

For each component and covariance direction, the relationship across genes was modelled as

$$E^C_{gakd} = \beta_{0akd} + \beta_{1akd} E^T_{gakd} + \epsilon_{gakd}. \tag{S13}$$

The slope $t$ statistic summarized whole-transcriptome correspondence. Positive values indicate concordant gene-expression profiles; negative values indicate inverse profiles. A large negative statistic is not absence of correspondence.

For an ordinary simple linear regression with an intercept and $G$ valid gene pairs, the algebraic relationship between the slope statistic and Pearson correlation is

$$t = r\sqrt{\frac{G-2}{1-r^2}}. \tag{S14}$$

The raw regression slope is not numerically equivalent to $r$ unless the variables have the appropriate scaling. A large nominal gene-wise statistic does not establish donor-level precision or overcome dependence among genes and spatial samples.

For each component, the covariance direction with the larger absolute slope statistic was retained for display, with its sign preserved (Figure 5 of the main manuscript). This direction selection was descriptive.

##### S8.4 Targeted core–matrix-associated markers

*PVALB* and *CALB1* encode parvalbumin and calbindin, respectively. They were selected as markers previously related to core–matrix-associated thalamic organization (Jones, 1998; Müller et al., 2020). Their analysis tests relative localization of selected molecular fields, complementing the cross-structure relationship across the whole transcriptome. It does not classify each component into a pure core or matrix cell population (Phillips et al., 2019).

For each marker $g$ and the dominant covariance direction $d^*_{ak}$, the component score was standardized across the algorithm–component comparison set $\mathcal{C}$:

$$z_{gak} = \frac{E^T_{gakd^*_{ak}} - \mathrm{mean}_{(a',k')\in\mathcal{C}}\, E^T_{ga'k'd^*_{a'k'}}}{\mathrm{sd}_{(a',k')\in\mathcal{C}}\, E^T_{ga'k'd^*_{a'k'}}}. \tag{S15}$$

Positive values indicate relatively high expression within this comparison set, not enrichment against an external spatial null. Standardization was performed separately for the two genes.

The AHBA is an adult post-mortem reference, whereas the PNC analytical sample spans childhood through young adulthood. These analyses characterize spatial molecular correspondence; they do not measure developmental expression changes in the imaged participants. Bulk-tissue marker expression does not directly determine cell-type proportions or projection anatomy.
